# Phertilizer: Growing a Clonal Tree from Ultra-low Coverage Single-cell DNA Sequencing of Tumors

**DOI:** 10.1101/2022.04.18.488655

**Authors:** Leah L. Weber, Chuanyi Zhang, Idoia Ochoa, Mohammed El-Kebir

## Abstract

Emerging ultra-low coverage single-cell DNA sequencing (scDNA-seq) technologies have enabled high resolution evolutionary studies of copy number aberrations (CNAs) within tumors. While these sequencing technologies are well suited for identifying CNAs due to the uniformity of sequencing coverage, the sparsity of coverage poses challenges for the study of single-nucleotide variants (SNVs). In order to maximize the utility of increasingly available ultra-low coverage scDNA-seq data and obtain a comprehensive understanding of tumor evolution, it is important to also analyze the evolution of SNVs from the same set of tumor cells.

We present Phertilizer, a method to infer a clonal tree from ultra-low coverage scDNA-seq data of a tumor. Based on a probabilistic model, our method recursively partitions the data by identifying key evolutionary events in the history of the tumor. We demonstrate the performance of Phertilizer on simulated data as well as on two real datasets, finding that Phertilizer effectively utilizes the copynumber signal inherent in the data to more accurately uncover clonal structure and genotypes compared to previous methods.

**Availability:** https://github.com/elkebir-group/phertilizer

## 1 Introduction

Cancer results from an evolutionary process that yields a heterogeneous tumor composed of multiple subpopulations of cells, or *clones*, with distinct sets of somatic mutations [1] (Fig. 1a). These mutations include single-nucleotide variants (SNVs) that alter a single base and copy-number aberrations (CNAs) that amplify or delete large genomic regions. Over the last decade, new developments in single-cell DNA sequencing (scDNA-seq) methods have helped uncover a wealth of insights regarding intra-tumor heterogeneity and cancer evolution [2–5]. In particular, the ongoing development and application of high-throughput, ultralow coverage scDNA-seq technologies (< 1×), such as direct library preparation (DLP+) [6] and acoustic cell tagmentation (ACT) [7], have paved the way for enriching our understanding of the role CNAs play in cancer progression and tumor evolution [6–8].

**Figure 1:**
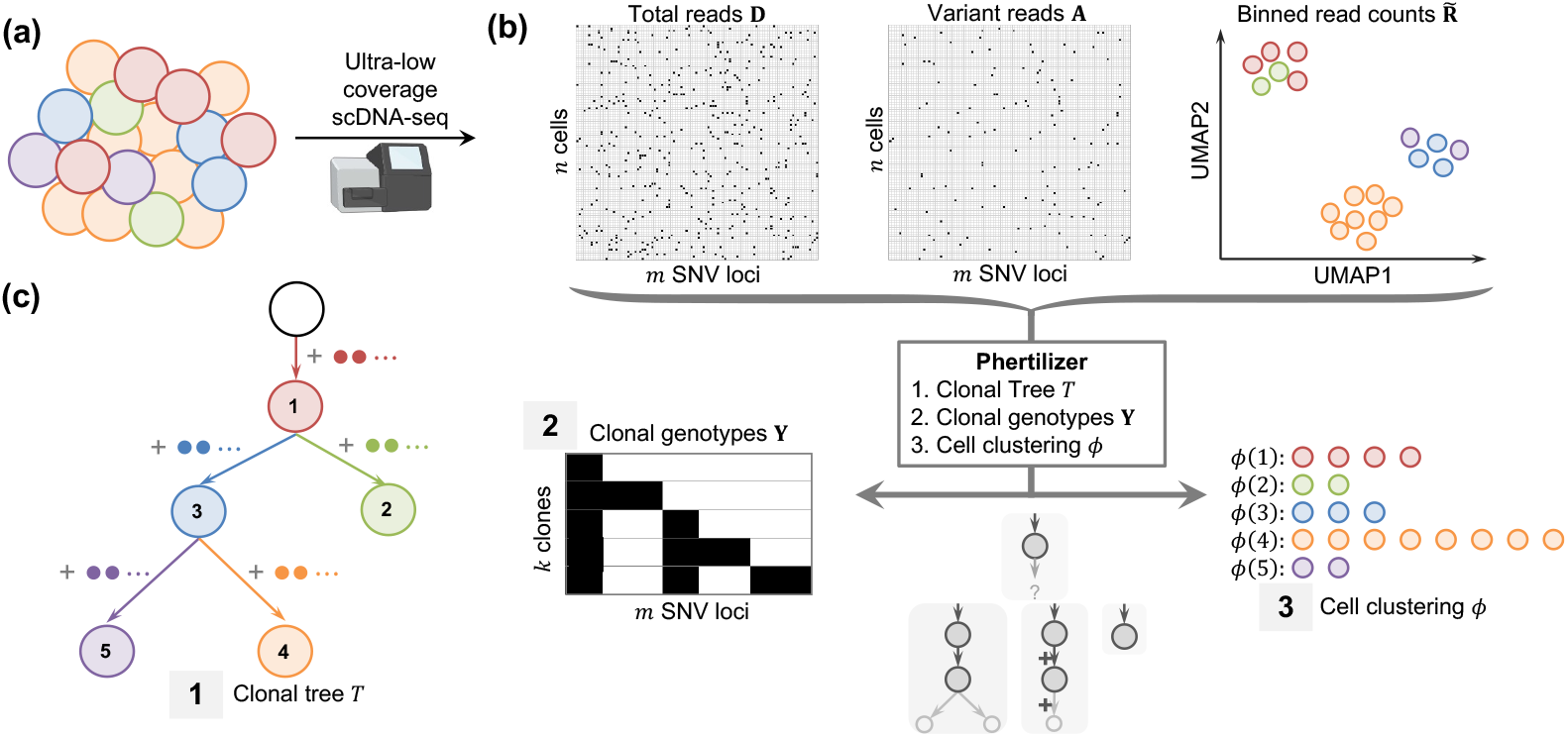
Phertilizer infers a clonal tree *T*, clonal genotypes Y and a cell clustering *ϕ* given ultralow coverage single-cell sequencing data. (a) A tumor consists of clones with distinct genotypes. (b) Ultra-low coverage scDNA-seq produces total read counts 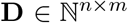 and variant read counts 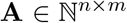 for *n* cells and *m* SNV loci, and low dimension embedding of binned read counts 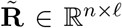. (c) Given maximum copy number *c* and sequencing error probability *α*, Phertilizer infers a clonal tree *T*, clonal genotypes **Y** and cell clustering *ϕ* with maximum posterior probability *P*(*T*, **Y**, *ϕ* | **A**, **D**, 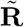, *c, α*).

The advantage these ultra-low coverage scDNA-seq technologies have over other high-throughput scDNA-seq methods (> 1×), like Mission Bio Tapestry [9], is the uniformity of coverage. This uniformity implies that the observed read counts for a genomic region is proportional to copy number, making it ideal for the analysis of subclonal CNAs that occur in only a small subset of tumor cells. However, this uniformity comes at the cost of sequencing depth, making it very difficult to identify and characterize the evolution of SNVs from ultra-low coverage scDNA-seq. Critically, to comprehensively study the evolution of a tumor from the same set of cells, both CNAs and SNVs should ideally be characterized by a single tumor phylogeny that depicts their coevolution. While this remains a long-term goal for the field, a first step in this direction is to increase our understanding of SNV evolution from ultra-low coverage scDNA-seq data by incorporating reliable copy number information into the inference of SNV tumor phylogenies.

Although phylogeny inference methods from bulk sequencing and cell clustering from single-cell RNA sequencing are expanding to incorporate both SNV and CNA features, such as TUSV-ext [10] and CASIC [11], current methods for tumor phylogeny and/or clone inference from single-cell sequencing naturally tend to focus on the features (SNV or CNA events) for which the data is ideally suited [12–26]. One exception in the medium to high coverage scDNA-seq regime is SCARLET [27], which refines a given copy number tree using SNV read counts under a CNA loss supported evolutionary model. While SCARLET accounts for sequencing errors and missing data, it was not designed to handle the extreme sparsity of ultralow coverage scDNA-seq. SBMClone [28] took the first step of using ultra-low coverage sequencing data to infer SNV clones via stochastic block modeling. Despite good performance on simulated data, especially with higher coverage (> 0.2×), SBMClone was unable to identify clear structure in a 10X Genomics breast cancer dataset [4] without *ad hoc* use of additional copy number clone information. Moreover, it is non-trivial to convert the inferred parameters of SBMClone’s stochastic block model to clonal genotypes, which may impact downstream analyses. Similarly, given a set of candidate SNV loci, SECEDO [29] first calls SNVs using a Bayesian filtering approach and then subsequently clusters cells using the called SNVs. While both of these clustering methods capitalize on the ever increasing throughput of ultra-low coverage scDNA-seq methods, neither method constrains the output by a tree and CNA features are used only in an *ad hoc* manner or for orthogonal validation. As a result, both methods imply that CNA and SNV data features should be segregated and analyzed in separate bioinformatics pipelines. The other emerging trend from the analysis of ultra-low coverage scDNA-seq is pseudobulk analysis [6]. This approach begins by identifying copy number clones using existing methods, followed by pooling cells that belong to the same CNA clones into pseudobulk samples, which are then independently analyzed to identify SNVs. Finally, phylogeny inference is performed with the copy number clones as the leaves of the tree. By doing so, this method does not allow for further refinement of these clones based on SNV evolution.

Here we introduce Phertilizer, the first method to infer an SNV clonal tree from ultra-low single-cell DNA sequencing of tumors. To overcome SNV coverage sparsity in this type of data, we leverage the strong copy number signal inherent in the data to guide clonal tree inference. By analogy to the planting and growing of trees, Phertilizer seeks to grow a clonal tree with maximum posterior probability by recursively inferring elementary clonal trees as building blocks. Our simulations demonstrate that Phertilizer accurately infers phylogenies and cell clusters when the number of cells matches current practice. In particular, Phertilizer outperforms a current method [28] for simultaneously clustering SNVs and cells as well as another commonly used *ad hoc* approach [6]. On real data, we find that Phertilizer effectively utilizes the copy-number signal inherent in the data to uncover clonal structure, yielding high-fidelity clonal genotypes.

## 2 Problem statement

Our goal is to infer an SNV phylogeny, guided by copy number aberrations, from ultra-low coverage sequencing data consisting of *n* cells and *m* identified single-nucleotide variants (SNVs). More precisely, we are given variant reads **A** = [*a_iq_*] and total reads **D** = [*d_iq_*], where *a_iq_* and *d_iq_* are the variant and total read counts for SNV locus *q* ∈ [*m*] in cell *i* ∈ [*n*], respectively (Fig. 1b). While the number of cells is large with the latest generation of ultra-low coverage single-cell DNA sequencing technology (*n* ≈ 1000 cells), the *coverage*, or the average number of reads that span a single locus is uniform but low (0.01 × to 0.5×). For example with a coverage of 0.01×, we would on average observe *a_iq_* = *d_iq_* = 0 reads for 99 out of every 100 loci q for each cell *i*. This sparsity renders phylogeny inference using SNVs extremely challenging. We propose to overcome this challenge using the following three key ideas.

First, similarly to current methods, we leverage the clonal structure present in tumors, i.e., cells typically cluster into a small number of clones. Thus, we seek to group the n observed cells into *k* clones (*k* ≪ n) via a cell clustering and corresponding clonal genotypes defined as follows:

### Definition 1.

Function *ϕ* : [*k*] → 2^[*n*]^ is a *cell clustering* provided its image encodes a partition of cells [*n*] into *k* (disjoint and non-empty) parts.

### Definition 2.

Matrix **Y** ∈ {0, 1}^*k×m*^ encodes *clonal genotypes* where *y_jq_* = 1 indicates that SNV *q* is present in clone *j* and *y_jq_* = 0 indicates that SNV *q* is absent in clone *j*.

Second, because ultra-low coverage scDNA-seq is sequenced uniformly, we may leverage the copy number signal inherent in the data to improve cell clustering performance [11] and guide tree inference. More specifically, we expect that all cells in a clone have identical copy number profiles. As we do not observe copy number directly, we will use observed reads counts 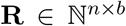, where *b* is the number of genomic bins, as a proxy for copy number. From read counts **R**, we derive distances that reflect copynumber similarity between pairs of cells on a low-dimensional embedding of binned read counts 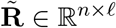 of **R** (i.e., *ℓ* ≪ *b*) — see Appendix A.1. Third, similarly to methods such as SCITE [12], SciCloneFit [15] and SPhyR [14], which operate on medium-to-high coverage scDNA-seq data, we consider that the observed cells are generated as the result of a tree-like evolutionary process that constrains the order of SNV clusters. In particular, we use the infinite sites model [30] defined as follows.

### Definition 3.

A tree *T* with nodes {*v*_1_,…,*v_k_*} rooted at node *v*_1_ is a *clonal tree* for clonal genotypes **Y** = [**y**_1_,…, **y**_*k*_]^T^ provided (i) each node *v_j_* is labeled by clonal genotype **y**_j_ and (ii) each SNV *q* is gained exactly once and subsequently never lost. That is, there exists no directed edge (*v_j′_*, *v_j_*) where the SNV is lost, i.e., *y_jq_* = 1 and *y_j′q_* = 0. Moreover, either the root node contains the SNV, i.e., *y*_1*q*_ = 1, or there exists exactly one directed edge (*v_j′_*, *v_j_*) where the SNV is introduced, i.e., *y_j′q_* = 0 and *y_jq_* = 1.

To relate the data to our latent variables of interest (T, **Y**, *ϕ*), we introduce a generative model in Appendix A.2 that describes the generation of variant read counts **A** and the binned read count embedding 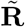. This model requires two hyperparameters *c* and *α*, where 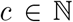 is the upper bound on the total number of chromosomal copies at any locus in the genome and *α* ∈ [0, 1] is the probability of misreading a single nucleotide during sequencing (Fig. S1). Importantly, while Definitions 1 to 3 explicitly indicate the number *k* of clones, the number *k* of clones is not a hyperparameter and will be part of the inference. Specifically, our generative model enables us to approximate the posterior probability *P*(*T*, **Y**, *ϕ* | **A**, **D**, 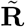, *c, α*) for a clonal tree T with any number k of nodes and associated clonal genotypes **Y** and cell clustering *ϕ* (derivation in Appendix A.2.4). However, due to limitations of the sequencing technology the number of clones that are detectable from the data may be fewer than the number of nodes in clonal tree *T*. The ability to detect clone *j* from the observed data is a function of the sequencing coverage, the number of cells in clone *j* and the number of SNVs newly introduced in clone *j*. To prevent overfitting of the data, the pre-specified detection threshold 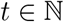 controls the minimum amount of observed data in support of each inferred clone. See Appendix A.3 and Fig. S2 for details and in particular Definition 4. This leads to the following problem.

### Problem 1

(Clonal Tree Inference (CTI)). Given variant reads 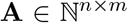, total reads 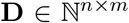, binned read count embedding 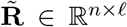, maximum copy number 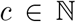, sequencing error probability *α* ∈ [0, 1] and detection threshold 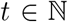, find a clonal tree *T* with detectable clonal genotypes **Y** and cell clustering *ϕ* with maximum posterior probability *P*(*T*, **Y**, *ϕ* | **A**, **D**, 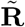, *c, α*).

## 3 Methods

To solve the CTI problem, Phertilizer maintains a set 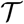 of candidate trees throughout three phases: (i) initialization, (ii) enumeration, and (iii) ranking each tree in 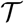 by its posterior probability. First, in the initialization phase, the set 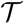 is initialized with a single tree containing only a root node *v*_1_. All n cells are assigned to the node’s cell cluster *ϕ*(*v*_1_) and all genotypes are initialized to *y*_*v*1,*q*_ = 1 for each SNV *q*.

Second, in the enumeration phase, Phertilizer recursively constructs the candidate set 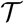 of clonal trees and the respective clonal genotypes and cell clusterings by performing three different elementary tree operations (`Linear, Branching` and `Identity`) on each leaf node *v_j_* of each candidate tree 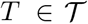 (Fig. 2). Specifically, each operation takes as input (*T*, **Y**, *ϕ*) and yields a new clonal tree *T′* with updated genotypes **Y′** and cell clustering *ϕ*′ by extending leaf *v_j_* of *T* (Fig. 2). The key idea is that each elementary tree operation breaks down the CTI problem into smaller subproblems. Intuitively, a `Linear` operation (Fig. 2a) replaces a leaf node with a two node linear subtree, while a `Branching` operation (Fig. 2b) replaces the leaf node with a three node binary subtree. The former represents stepwise acquisition of SNVs with evidence of intermediary clones present in the observed data, while the latter indicates evidence of divergence from a common ancestor [31]. The `Identity` (Fig. 2c) operation does not modify the tree.

**Figure 2:**
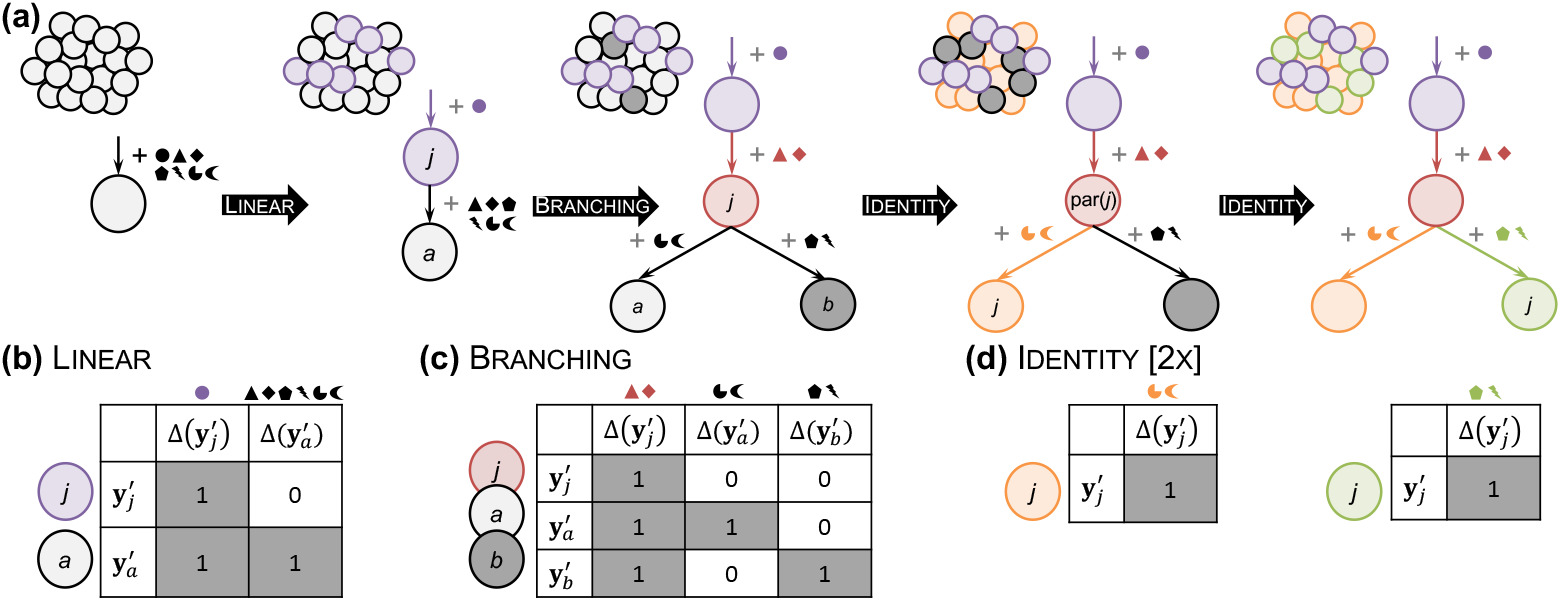
Phertilizer solves the CTI problem by enumerating clonal trees using three elementary operations in a recursive fashion. (a) Each operation yields a new clonal tree *T′* by extending a leaf *v_j_* of the previous clonal tree *T* and reassigning its SNVs Δ(y_*j*_) and cells *ϕ*(*j*). The resulting clonal genotypes **Y**′ and cell clustering *ϕ′* are constrained as depicted: (b) `Linear`, (c) `Branching`, and (d) `Identity`.

These operations are defined more formally in Appendix A.4. While the specifics vary slightly, both `Linear` (Appendix A.4.1) and `Branching` (Appendix A.4.2) are solved using a coordinate descent approach. That is, we fix clonal genotypes **Y**′ and solve for the cell clustering *ϕ′* and alternate. Drawing parallels between image segmentation, where combining both pixel and pixel location features results in better clustering [32], we incorporate the binned read count embedding 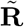 and variant read counts **A** into a single feature [11]. We then use this feature as input to the normalized cut algorithm [32] to obtain a cell clustering with two clusters. The advantage of combining SNV and CNA signal into a single feature is that cell clustering is improved when one or both of these signals are weak, subsequently improving the SNV partition which we solve for next. Given a fixed cell clustering *ϕ′*, we use our generative model to update clonal genotypes **Y′** by assigning each SNV to the node in extended tree *T′* with maximum posterior probability. We terminate this process upon convergence or when a maximum number of iterations is reached. The resulting clonal tree *T′* is then appended to the candidate set 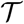 provided all its clone are detectable for a specified detection threshold *t* (Appendix A.3) and meet additional regularization criteria (Appendix A.4.3).

Third, once no new clonal trees are added to the candidate set 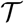, we return the clonal tree *T*, clonal genotypes **Y** and cell clustering *ϕ* with maximum posterior probability after post-processing (Appendix A.4.4). Importantly, the top down approach by Phertilizer requires that no assumptions be made *a priori* regarding the number of nodes *k* in the inferred clonal tree. Phertilizer is implemented in Python 3, open source (BSD-3-Clause), and available at https://github.com/elkebir-group/Phertilizer.

## 4 Results

### 4.1 Simulation study

#### Overview

To assess the performance of Phertilizer and compare it with previously proposed methods, we performed a simulation study with known ground truth clonal trees, evaluating the following four questions: (i) How accurate are the inferred clonal trees? (ii) How well is each method able to identify clusters of cells with similar clonal genotypes? (iii) How accurate are the inferred clonal genotypes? (iv) How sensitive is each method to violations of the infinite sites assumption? We designed our simulation study to match the characteristics of current datasets generated from ultra-low coverage scDNA-seq. To achieve this, we generated simulation instances with varying number *k* ∈ {5, 9} of nodes, number *n* ∈ {1000, 2000} of sequenced cells and number *m* ∈ {5000, 10000, 15000} of SNVs with a mean sequencing coverage *g* ∈ {0.01 ×, 0.05 ×, 0.1 × }. We replicated each of these combinations 10 times for a total of 360 instances. See Appendix B.1 for details on the simulation instances.

We assessed the quality of an inferred solution (*T*, *ϕ*, **Y**) against a ground-truth tree *T*\*, cell clustering *ϕ*\* and clonal genotypes **Y**\* using ancestral pair recall (APR), incomparable pair recall (IPR), and clustered pair recall (CPR) metrics for cells and SNVs [33], as well as genotype similarity. In addition, we computed a single *accuracy* value (domain: [0, 1]) composed of the weighted average of APR, IPR and CPR — where the weights are proportional to the number of pairs in each class — such that an SNV and cell accuracy of 1 imply that the inferred solution perfectly matches ground truth. The *genotype similarity* equals 1 minus the normalized Hamming distance between ground truth genotypes and inferred genotypes of cells. We refer to Fig. 3a and S6 for examples and to Appendix B.1.4 for more formal definitions.

**Figure 3:**
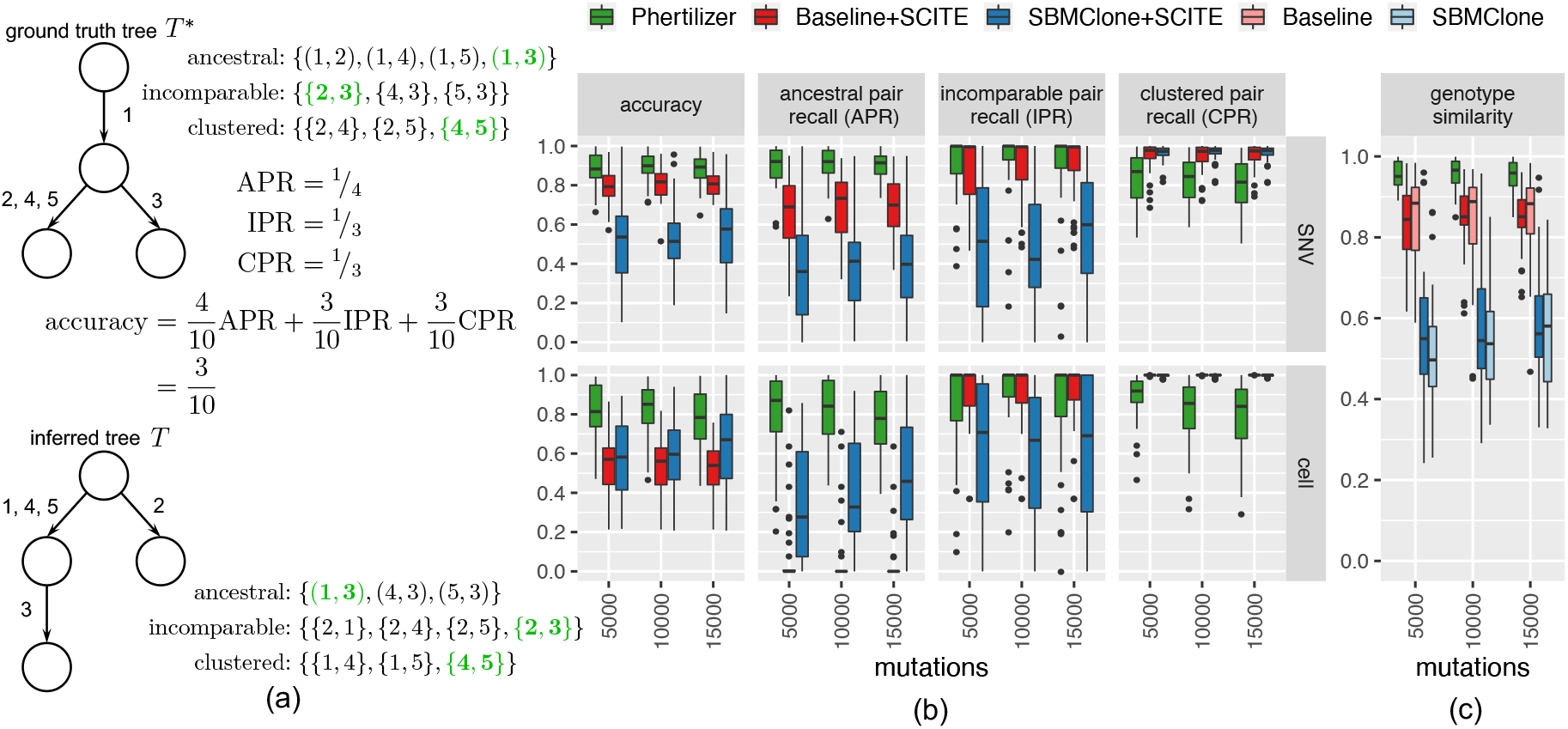
Phertilizer outperforms Baseline and SBMClone on simulated data. We show aggregated results for *n* ∈ {1000, 2000} cells, *k* ∈ {5, 9} clones and coverage *g* = 0.05×. (a) Example of SNV accuracy, ancestral pair-recall (APR), clustered pair recall (CPR), and incomparable pair recall (IPR) metrics. See Fig. S6 for the corresponding example for cell metrics and genotype similarity. (b) Phertilizer outperforms Baseline +SCITE and SBMClone +SCITE in APR and IPR for both SNVs and cells. Although competing methods rank higher in CPR, overall Phertilizer performs the best considering the accuracy. (c) Phertilizer more accurately recovers clonal genotypes than competing methods.

We benchmarked against SBMClone [28] because it is the only existing clustering/genotyping method that co-clusters cells and SNVs. We also benchmarked against a commonly adopted *ad hoc* practice which we refer to as Baseline [6, 7]. In Baseline, cells are first clustered into clones from the read count embedding 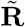. Clonal genotypes are obtained by pooling the reads of all cells assigned to clone *j* and setting *y_jq_* = 1 when the ratio of alternate to total reads at each locus *q* exceeds a threshold of 0.05 (details provided in Appendix B.1.1). Since there exists no standalone method that performs cell clustering, genotyping and tree inference for ultra-low coverage scDNA-seq, we paired SBMClone and Baseline with SCITE [12].

For brevity, we focus our discussion on a coverage *g* = 0.05× and show aggregated results for *n* ∈ {1000, 2000} cells and *k* ∈ {5, 9} clones. We report the median for all performance metrics and include the interquartile range (IQR), i.e., the difference between the 75th and 25th percentiles of the data, when appropriate. We note deviations in trends where relevant, and refer to Appendix B for remaining results.

#### Results

We first evaluated the accuracy of SNV placement on the inferred tree by assessing APR, CPR, IPR and accuracy for SNVs (Fig. 3b, Fig. S8). Overall, Phertilizer achieved the highest SNV accuracy (median: 0.90) in terms of SNV placement of all three methods (Baseline+SCITE median: 0.81, SBM-Clone+SCITE median: 0.54). For SNV APR, Phertilizer (median: 0.92) outperformed both Base-line+SCITE (median: 0.71) and SBMClone+SCITE (median: 0.38). This implies that our `Linear` operation reliably partitions SNVs accurately and that our `Branching` operation performs well at identifying SNVs that should be placed at the parent node. Moving on to SNV IPR, Phertilizer (median: 1.0, IQR: 0.11) outperformed SBMClone+SCITE (median: 0.48, IQR: 0.5) with lower variability than Baseline+SCITE (median: 1.0, IQR: 0.18). Thus, in addition to correctly identifying the SNVs in the parent node, the `Branching` operation also successfully partitions the SNVs in the children nodes. However, for SNV CPR, Phertilizer performed worse compared to SBMClone+SCITE (median: 0.98) and Baseline+SCITE (median: 0.97) but still maintained good performance (median 0.84). It is important to note that an SNV CPR of 1 is achievable by clustering all SNVs into a single cluster. Thus, the cost of underfitting the data, or grouping the SNVs into a few very large clusters, will be reflected in decreased APR and IPR. Indeed, we observed this to be the case for both Baseline+SCITE, which performed relatively poorly on APR, and SBMClone+SCITE which was the worst performing on both APR and IPR.

Next, we evaluated the accuracy of the cell clustering and placement on the inferred tree. We observed similar trends across accuracy, APR, IPR and CPR cell performance metrics to those identified for their SNV counterparts (Fig. 3b, Fig. S8). Phertilizer achieved the highest overall cell accuracy (median: 0.82) in comparison to SBMClone+SCITE (median: 0.60) and Baseline+SCITE (median: 0.56). Likewise, Phertilizer (median: 0.84) outperformed all other methods on cell APR (Baseline+SCITE median: 0.0, SBMClone+SCITE median: 0.32). On cell IPR, both Phertilizer (median: 1.0, IQR: 0.2) and Baseline+SCITE (median: 1.0, IQR: 0.13) significantly outperformed SBMClone+SCITE (median: 0.67, IQR: 0.68) but Baseline+SCITE had slightly lower variability (IQR) than Phertilizer. Similarly to SNV CPR, Baseline+SCITE (median: 1.0) and SBMClone+SCITE (median: 1.0) outperformed Phertilizer on cell CPR (median: 0.87), but the corresponding decreased performance in cell APR and IPR was indicative of inferring too few cell clusters. In addition to providing more supporting evidence for the validity of our elementary operations, these cell placement performance metrics also highlight the advantage of Phertilizer utilizing both copy number and SNV signal for tree inference. In contrast, Baseline+SCITE prioritizes copy number signal and is unable to further refine a cluster of cells with distinct clonal genotypes but the same copy number profile. Conversely, SBMClone+SCITE ignores copy number signal and struggles to infer clones with sparse SNV signal.

Finally, we assessed the genotype similarity and included SBMClone and Baseline as this is obtained prior to tree inference. Since genotype similarity compares the inferred genotype or each simulated cell against its ground truth genotype, it captures the interplay between cell clustering and clonal genotyping. Given that Phertilizer achieved the highest performance on the clonal tree inference and cell clustering metrics, we would expect Phertilizer to have the highest genotype similarity. Indeed, Fig. 3c demonstrates that to be true since Phertilizer was the only method to have a median genotype similarity above 0.95. Baseline was the next highest (median: 0.88), closely followed by Baseline+SCITE (median: 0.85), while SBMClone+SCITE (median: 0.55) and SBMClone (median: 0.53) had the worst performance. When evaluating these metrics at the lowest coverage *g* = 0.01×, Phertilizer maintained top performance on both cell placement and genotype similarity but Baseline+SCITE was competitive with Phertilizer on SNV placement (Fig. S7). For the highest coverage *g* = 0.1×, Phertilizer achieved the highest accuracy on SNV placement and cell placement and had a median genotype similarity of 0.97 while the next closest competitor (Baseline) had a median similarity of 0.89 (Fig. S9). In terms of running time for the case of n = 1000 cells, m = 15000 SNVs, and a coverage of g = 0.01×, the median running time of Phertilizer was 460 s, 45.9 s for Baseline+SCITE and 101 s for SBMClone+SCITE (Fig. S12).

To perform sensitivity analysis, we generated two additional sets of simulations. The first had the same parameters as above but excluded CNAs, such that every locus was heterozygous diploid. We also excluded Baseline+SCITE from comparison as only a single clone was inferred after cell clustering. We found Phertilizer still outperformed SBMClone+SCITE but that for coverage *g* = 0.01× our performance were slightly worse than simulations with CNAs (Fig. S10), implying CNA features aid inference when sequencing coverage is extremely sparse. For the second, simulations were generated under a Dollo [34] evolutionary model with *k* = 9, *m* = 15000, coverage *g* ∈ {0.01×, 0.05×, 0.01×}. We found that Phertilizer still outperformed Baseline+SCITE and SBMClone+SCITE, having maintained high scores on all performance metrics (Fig. S11) with the exception of cell APR for the lowest coverage (Fig. S11a).

In summary, we conclude that given the high level of accuracy obtained by Phertilizer on these performance metrics, not only are the elementary operations successful in isolation but also the posterior probability is useful in discriminating between candidate clonal trees. Additionally, we find that utilizing copy number signal, whenever available, is necessary when working with ultra-low coverage scDNA-seq data but not sufficient for accurate clonal tree reconstruction and/or SNV genotyping.

### 4.2 High-grade serous ovarian cancer patient

Utilizing a grid search over input parameters(Appendix B.2.1), we ran Phertilizer on *n* = 890 DLP+ sequenced cells from three clonally-related cancer cells lines sourced from the same high-grade serous ovarian cancer patient [6]. We used the variant and total read counts **A**, **D** for *m* = 14,068 SNVs and derived binned read counts **R** for *b* = 6,207 bins (bin width of 500 KB) from data reported by Laks et al. [6]. The average sequencing coverage for these data was 0.25×. Utilizing an approach similar to the Baseline method described above, Laks et al. [6] identified 9 copy number clones (labeled A-I) via dimensionality reduction and density-based clustering and reconciled them in a phylogeny with the copy number clones as leaves (Fig. 4a). We annotated inferred trees with mutations in cancer-related genes in the cBioPortal [35, 36] and Cancer Gene Census (CGC) [37] from COSMIC v97 (see Appendix B.2.2).

**Figure 4:**
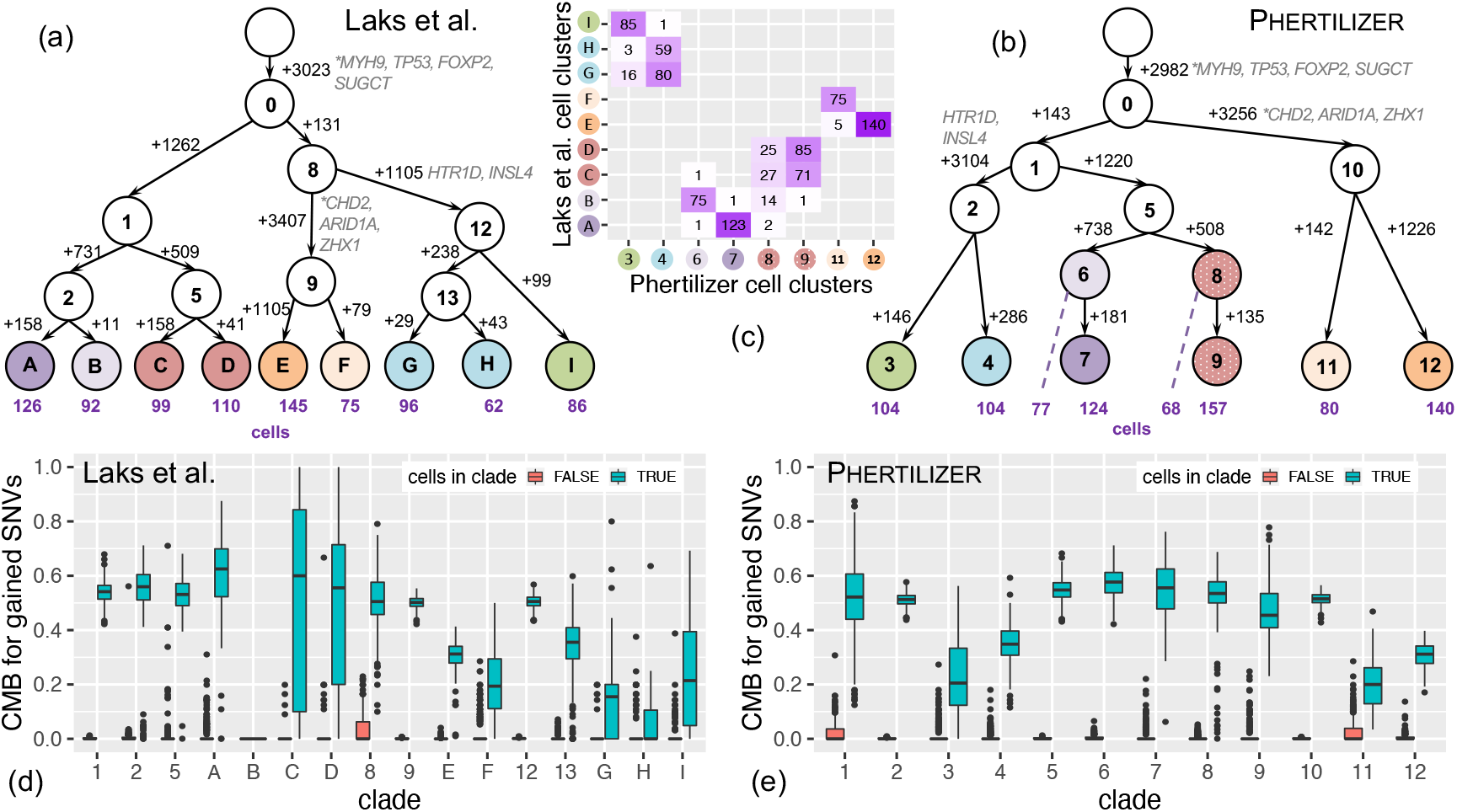
Phertilizer improves upon the SNV phylogeny previously inferred by Laks et al. [6]. (a) The clonal tree inferred by Laks et al. [6] with edges labeled by SNV gains and cell numbers shown below leaf nodes. (b) The clonal tree inferred by Phertilizer with edges labeled by SNV gains and cell numbers shown below leaf nodes. Cancer-related genes are labeled next to the SNVs in (a) and (b) with ‘*’ indicating a stop-gain variant. (c) Mapping between the Laks et al. [6] cell placement and Phertilizer cell placement. (d-e) Cell mutational burden (CMB) comparison between cells within (blue) and outside (red) of each clade in the inferred Laks et al. [6] and Phertilizer clonal tree.

As shown in Fig. 4b, Phertilizer inferred a clonal tree with 13 nodes and 8 clones. We found that Phertilizer’s tree closely aligned with the Laks et al. [6] tree, with both approaches correctly identifying three major clades corresponding to the three distinct cell lines of origin (Fig. 4c). Additionally, driver genes *TP53, SUGCT*, and *MYH9* are identified as clonal in both the Laks et.al [6] inferred tree (Fig 4a) and the Phertilizer clonal tree (Fig 4b). Similarly, subclonal SNVs in *CHD2, ARID1A, ZHX1, HTR1D*, and *INSL4* were placed in the corresponding clades of both trees. To further assess the quality of each inferred clade, we developed a performance metric called *cell mutational burden* (CMB) defined as CMB(*i, M*) = ∑_*q*∈*M*_ **1**{*a_iq_* > 0}/ ∑_*q*∈*M*_ **1**{*d_iq_* > 0}. In words, CMB(*i, M*) is the fraction of mapped SNV loci *M* with mapped variant reads in cell *i*. For a specified *clade j* or subtree rooted at node *v_j_*, SNVs *M_j_* are the SNVs gained at node *v_j_*. For a cell *i* placed within clade *j*, we expect CMB(*i, M_j_*) to be high, although the value will depend on copy number. By contrast, for cells placed outside of clade *j*, we expect CMB(*i, M_j_*) to be low. More details on CMB are provided in Appendix B.2.3 and Fig. S13.

Fig. 4d,e depicts the comparison of the distributions of CMB for all clades for cells placed within and outside that clade for Laks et al. [6] and Phertilizer, respectively. For cells placed outside of a specified clade, the reported tree by Laks et al. [6] as well as the tree inferred by Phertilizer have a median CMB of 0 for all clades. This is indicative of high specificity of both methods, i.e., SNVs are not assigned to a clade if there are observations of that SNV outside that clade. However, for cells placed within the clades, we observed greater variability for the Laks et al. [6] inferred clades than Phertilizer. This variability was most pronounced for the leaf nodes, especially C, D, G, F, *H* and *I*, where most have a small number of SNVs gained. We further analyzed clusters G and H, where the 25th percentile of the CMB for cells within the clade equals 0, and clusters C and D that had large variability (IQR of 0.74 and 0.51, respectively).

The location of clusters G and H in the embedding space suggests overfitting during density-based clustering, with an arbitrary split of a larger cohesive cluster containing G and H (Fig. S14a). In comparison, Phertilizer uses both copy number and SNV signal, resulting in the Laks et al. [6] G and H cells being clustered together to node 4 in the inferred Phertilizer clonal tree. Comparing CMB distributions for cells within clades G and H (Fig. 4d) with Phertilizer’s inferred clade 4 (Fig. 4e), we observed a higher 25th percentile (0.31) for node 4 than for G (0.00) and H (0.00). This resulted in large separation between the CMB distributions of cells within and outside of clade 4 for Phertilizer but not for Laks et al. [6] G and H clades, implying better SNV placement in Phertilizer’s clonal tree.

The last major difference of note between the two inferred trees is the clustering of the cells in Laks et al. [6] nodes C and D versus Phertilizer’s nodes 8 and 9. Similarly to nodes G and H, we did not observe a clear separation of C and D in the embedding space (Fig. S14a), making it difficult to define clusters of these cells without more rigorous copy number profiling and clustering. However, as we saw in the simulation study, Phertilizer was able to detect further SNV evolution within a set of cells having the same copy number profile. Cells in these clusters would not be split in half in the embedding space but instead should be scattered randomly throughout a single cluster in the embedding space. In addition to having observed cells in nodes 8 and 9 randomly scattered in a cluster in the embedding space (Fig. S14b), we also observed a clear separation between the CMB distributions for cells placed within and outside of clades 8 and 9 (Fig. 4e). For these clades, we also observed low variability with IQRs of 0.08 and 0.13, respectively, whereas clusters *C* and *D* have very high variability in the inferred Laks et al. [6] clonal tree.

Overall, both inferred clonal trees are very similar but since Phertilizer simultaneously uses both CNA and SNV information, we obtained a slight improvement in terms of SNV phylogeny inference. Additionally, we note that the small number of SNVs gained at most of the leaf nodes is a direct result of the bottom up approach taken by Laks et al. [6], which performed pseudobulk SNV calling individually on each cell cluster. When SNVs are present in multiple cell clusters but at low prevalence in each cluster, they may not pass filtering of current somatic SNV callers for a cell cluster. This gives the appearance that SNVs are unique to a single clone, when in actuality they are present in multiple clones but have not been called. When sequencing coverage is closer to the 0.01× as opposed to the 0.25× that we have with these data, correctly genotyping a cell cluster becomes more challenging. In contrast, the top down approach of Phertilizer is better suited to detect SNVs present in multiple cell clusters at low prevalence.

### 4.3 Eight triple negative breast tumors

We applied Phertilizer to eight triple negative breast tumors sequenced via ACT [7], labeled TN1 to TN8. After dimensionality reduction of normalized and GC-bias corrected binned read counts **R**, Minussi et al. [7] identified for each tumor two sets of cell clusters with varying granularity, denoted as superclones and subclones. To obtain the input set of SNVs for each patient, we performed SNV calling of a pseudobulk sample of pooled sequenced cells using MuTect2 in tumor-only mode [38]. See Appendix B.2.5 for further details on data processing. Table S1 displays the breakdown of each tumor in terms of the number n of cells, the number m of SNVs and average coverage g, and depicts the number of inferred clones by Phertilizer, Minussi et al. [7] superclones and subclones, as well as the number of inferred clones by SBMClone. Note that these data have a markedly lower coverage (ranging from 0.017× to 0.039×) than the DLP+ data (0.25×). We additionally ran Baseline+SCITE with the cell clusters fixed to the Minussi et al. [7] subclones but all instances except TN3 and TN5 timed out after 10 hours. However, the CMB distribution for the inferred trees for TN3 and TN5 (Fig. S16) provides no evidence in support of these trees. SBMClone only inferred a single clone for all patients, while Phertilizer inferred a tree with more than one clone for 4 out of the 8 tumors (TN1: 6, TN2: 6, TN4: 4 and TN8: 2). These four tumors have the highest average coverage of the eight patients. We will focus our discussion on the clonal trees inferred by Phertilizer for tumors TN1 (Fig. 5) and TN2 (Fig. S17).

**Figure 5:**
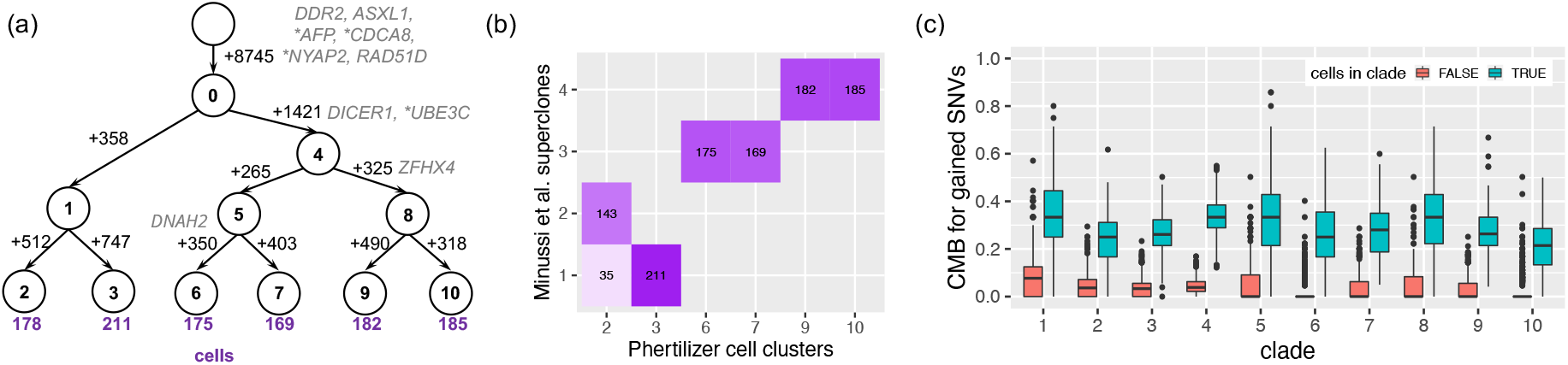
Phertilizer infers clonal tree for breast cancer tumor TN1. (a) The tree inferred by Phertilizer with numbers of SNVs labeled beside the edges, and numbers of cells labeled beneath the leaves. Cancer-related genes are labeled next to the SNVs (‘*’: stop-gain variant). (b) A mapping between Phertilizer’s cell clusters and the Minussi et al. [7] superclones. (c) The cell mutational burden (CMB) comparison between cells within (blue) and outside of (red) each clade in the inferred clonal tree.

For tumor TN1, Phertilizer inferred a branching tree with 11 nodes and 6 clones (Fig. 5a). We also identified a subclonal missense SNV in driver gene *DICER1* associated with tumorigenesis and poor prognosis [39,40]. We noted good concordance between the Minussi et al. [7] superclones and the Phertilizer cell clusters, with the exception of 35 cells that appear as outliers in superclone 1 (Fig. 5b, Fig. S15). This suggests these cells might fit better in superclone 2 based on SNV signal. In addition, we identified 8745 of the 13934 SNVs as truncal. This large truncal distance and branching structure were in alignment with the truncal distance in the clonal lineage tree inferred by Minussi et al. [7] using bulk whole exome sequencing. We used CMB to assess the performance of SNV and cell placement (Fig. 5c). For clades 5 through 10, we observed that the median CMB for cells outside of the clade is 0. Clades 9 and 10 were particularly interesting because the embedding space depicts the occurrence of SNV evolution within Minussi et. al.’s [7] superclone 4 (Fig. S15). For clades 2 through 4, we noted the median CMB for cells outside of the clade was around 0.035 for each of these clades, while clade 1 is the highest at 0.077. Upon further investigation of these 358 SNVs, we found that the median number of mapped reads was 5 when aggregating all cells, making these SNVs especially challenging to place. This drop in performance on the median CMB for cells outside of a clade when compared to the ovarian cancer patient (Section 4.2) is expected due to the drop in sequencing coverage from 0.25× to 0.031×. However, we still observed a large separation between CMB distributions for the cells within the clade and cells outside the clade for all clades.

For tumor TN2, we inferred a branching clonal tree with 11 nodes and 6 clones (Fig. S17). Two of Phertilizer’s cell clusters directly agreed with Minussi et al. [7] superclones. However, Phertilizer split the remaining two superclones into four cell clusters (3, 4, 5, 6) using SNV information. We observed low median CMB (0) for cells outside of the clade and a distinct separation between the cells within and outside of the clade distributions, providing evidence for this cell clustering and SNV placement. For tumor TN4, we identified an 8-node tree with five cell clusters, with trends similar to tumors TN1 and TN2 in terms of cell clustering concordance and CMB (Fig. S18). Finally, for tumor TN8 at the lowest coverage 0.021× of these four tumors, Phertilizer only inferred a 3-node branching tree with two cell clusters (Fig. S19).

## 5 Discussion

Ultra-low coverage scDNA-seq has greatly enhanced our ability to study tumor evolution from a copy number perspective [7, 17]. Capitalizing on the strong copy number signal inherent in this data, we proposed a new method, Phertilizer, that grows an SNV phylogeny in a recursive fashion using elementary tree operations. We demonstrated the effectiveness of our approach relative to existing clustering methods on both simulated and real data. Importantly, we found that for the current number of cells (800 – 2000) used in practice, Phertilizer performs markedly better than these methods, yielding more accurate clonal trees, cell clusters, and clonal genotypes. As the first method to reconstruct the evolutionary history of SNVs from ultra-low coverage scDNA-seq, Phertilizer helps advance the study of tumor evolution and makes progress towards the goal of joint SNV and CNA phylogeny inference at single-cell resolution.

There are several additional limitations and directions for future research. First, as sequencing coverage drops below 0.02× as in the ACT data, Phertilizer does not infer clonal trees with more than one clone. Although inference is impacted by numerous factors, like copy number profile, it does perhaps suggest a coverage of approximately 0.02× as the limit of detection for Phertilizer. Second, accurate variant calling also remains an open problem for ultra-low scDNA-seq data, making it challenging to identify input subclonal variants. Third, the infinite sites model used in this work is often violated due to copy-number deletions. While we demonstrate robustness to such violations, a future direction is to use the Dollo evolutionary model [14, 34]. Fourth, our model lacks an explicit placement of CNA events on the tree. Tree reconciliation methods, such as PACTION [41], can now be applied to integrate an SNV clonal tree generated by Phertilizer and a CNA tree to obtain a joint tree. Finally, beyond SNVs and CNAs, we plan to support structural variants and integrate other omics modalities such as methylation and transcription [42].

## Acknowledgements

We thank the Navin Lab and Darlan Minussi for their assistance with the ACT data. Additionally, we thank the Shah Lab, including Daniel Lai, Robert Reinert, and Andrew McPherson, for their assistance with the DLP+ data. M.E-K. was supported by the National Science Foundation (CCF-2046488) as well as funding from the Cancer Center at Illinois. I.O. was supported by a Gipuzkoa Fellows grant from the Basque Government, a Ramon y Cajal Grant from Spain, and a grant from the Spanish Ministry of Science and Innovation (PID2021-126718OA-I00). This work used resources, services, and support provided via the Greg Gulick Honorary Research Award Opportunity supported by a gift from Amazon Web Services.

---

**Figure S1:**
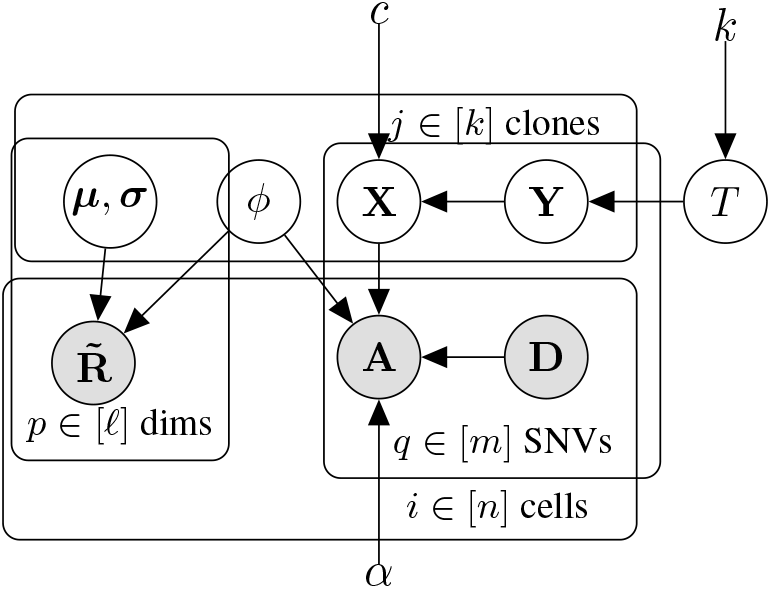
Plate diagram. The observed data consists of variant and total read counts **A**, **D** and binned read counts 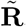, generated from an unobserved cell clustering *ϕ* and clonal tree *T* of *k* clones with clonal genotypes **Y**. Given hyperparameters *θ* = (*c, α*) and observed data (**A**, **D**, 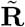), we seek (*T*, **Y**, *ϕ*) with maximum posterior probability *P*(*T*, **Y**, *ϕ*, | **A**, **D**, 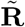, *θ*), marginalizing over latent variables **X**, *μ* and *σ*.

## A Supplementary methods

### A.1 Data processing

Phertilizer either takes as input binned read counts 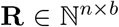 of *n* cells with read counts grouped into *b* bins or alternatively a binned read count embedding 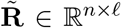 of read counts of *n* cells projected into *ℓ* ≪ *b* dimensional space. If provided the former, Phertilizer will project the binned read counts **R** into *ℓ* = 2 dimensions as follows. First, we compute normalized binned read counts 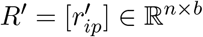 as

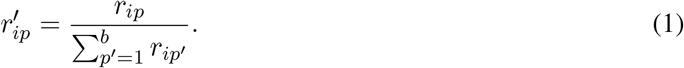

Next, we use UMAP [43] to project the normalized binned read counts **R′** to *ℓ* = 2 dimensions with the following parameter settings:

~~~
n_neighbors=40, spread=1, n_components=2, min_dist=0.
~~~

This yields the binned read count embedding 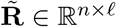 where *ℓ* = 2.

### A.2 Generative model

Our generative model (Fig. S1) captures the evolution and the subsequent process of ultra-low coverage sequencing of *n* cells with *m* SNVs yielding our data **A**, 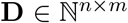 and 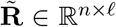. For a number 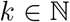 of clones, we generate a rooted tree *T* with *k* nodes with a uniform prior *P*(*T* | *k*). We generate clonal genotypes **Y** for tree *T* under the infinite sites model with *P* (**Y** | *T*) uniformly distributed. We model the cell clustering *ϕ* : [*k*] → 2^[*n*]^ with *P*(*ϕ*) distributed uniformly over all possible partitions of *n* cells into *k* clones/parts. The prior probability *P*(**D**) of total read counts **D** is also uniform.

The generative model has three remaining components: the (i) latent variant allele frequency (VAF) model, (ii) the variant read count model and (iii) the binned read count embedding model. First, the latent VAFs 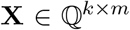 are dependent on the clonal genotypes **Y**. Thus, whenever clonal genotype *y_jq_* = 0 for SNV *q* of clone *j*, the latent VAF *x_jq_* = 0. Otherwise, *x_jq_* is the ratio of chromosomal copies that harbor SNV *q* to the total number of chromosomal copies of clone *j*. Appendix A.2.1 contains the precise definition of *P*(**X** | **Y**, *θ*). Second, given latent VAFs **X**, total read counts **D**, cell clustering *ϕ* and the probability *α* of misreading a single nucleotide during sequencing, we model variant read counts with a binomial distribution that accounts for sequencing errors. Appendix A.2.2 provides the derivation of *P*(**A** | **D**, **X**, *ϕ, θ*). Third, we model the generation of the binned read counts embedding 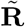 using a Gaussian mixture model where 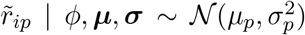. Latent variables 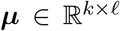 and 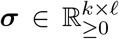 are independent for each clone *j* ∈ [*k*] and dimension *p* ∈ [*ℓ*]. Appendix A.2.3 contains the definition of 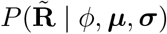.

This model requires two hyperparameters *c* and *α*, where 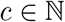 is the upper bound on the total number of chromosomal copies at any locus in the genome and *α* ∈ [0, 1] is the probability of misreading a single nucleotide during sequencing (Fig. S1). Given hyperparameters *θ* = (*c, α*) and observed data (**A**, **D**, 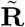), we seek to identify latent variables (*T*, **Y**, *ϕ*) with maximum posterior probability 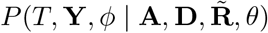. We show in Appendix A.2.4 that this can be approximated as 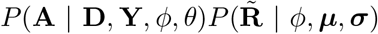 taking **μ** and **σ** as the maximum *a posteriori* estimates.

#### A.2.1 Latent variant allele frequency model

The latent variant allele frequency (VAF) *x_jq_* denotes the fraction of chromosomal copies that harbor the SNV at locus *q* for clone *j*. Let 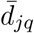 be the number of chromosomal copies of the SNV locus *q* in clone *j*, and let 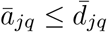 be the number of chromosomal copies that harbor the SNV. Mathematically, we define the latent VAF as 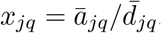.

The latent VAF 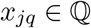 is conditioned on the latent clonal genotype *y_jq_* ∈ {0, 1}. Given a pre-specified maximum copy number 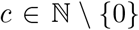, we define the multiset *S* of latent VAFs associated with SNV state *y_jq_* = 1 as

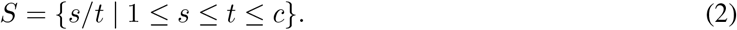

We denote with |{*x* ∈ *S*}| the number of occurrences of latent VAF *x* in the multiset *S*. Then, we have

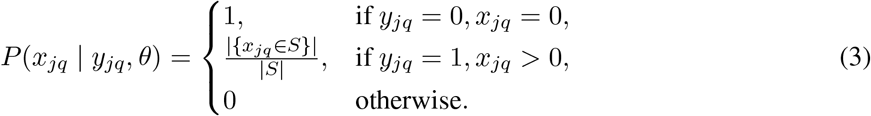

For example, if we have maximum copy number *c* = 2, the SNV is present (i.e., *y_jq_* = 1), we have that *S* = {1/1, 1/2, 2/2} and that *P*(*x_jq_* = 1/2 | *y_jq_* = 1,θ) = 1/3.

Without assuming any prior information on the chromosomal copy numbers in each clone *j*, this model assumes all possible combinations of 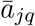 and 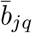 are equally likely. Using independence among clones [*k*] and SNV loci [*m*] when given clonal genotypes **Y**, we have that

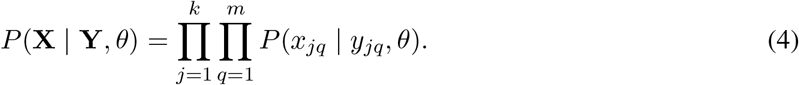

#### A.2.2 Variant read count model

Given latent VAFs **X**, total read counts **D** and cell clustering *ϕ*, we model alternate read counts **A** as a binomial distribution accounting for sequencing errors. While previous models for medium-to-high coverage scDNA-seq data specifically account for errors introduced during PCR amplification [44, 45], the lack of preamplification in the latest generation of ultra-low scDNA-seq technology restricts errors to base calling errors that occur during sequencing [6, 7]. Let *α* be the probability of misreading a single nucleotide during sequencing, next we derive *P*(**A** | **D**, **X**, *ϕ*, *θ*).

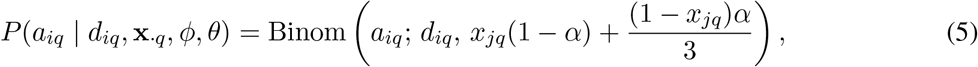

where *j* is the unique clone corresponding to cell *i*, i.e., *i* ∈ *ϕ*(*j*). Using an indicator variable, this is equivalent to

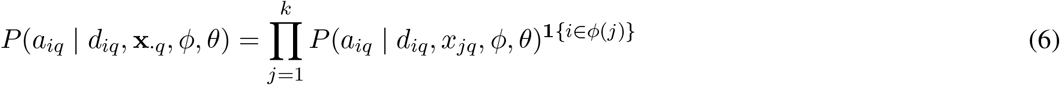

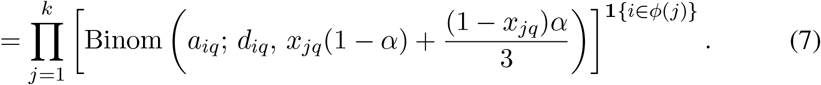

Per the graphical model, we have independence among cells [*n*] and SNV loci [*m*] when conditioning on **D**, **X**, *ϕ* and *θ*, yielding

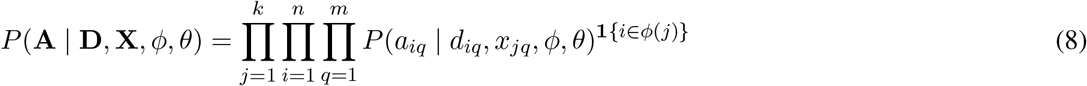

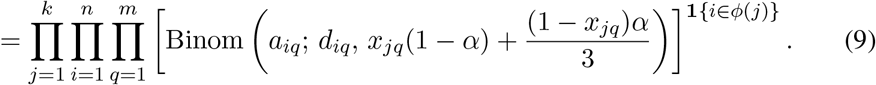

#### A.2.3 Binned read count embedding model

The projection of binned reads counts **R** into a low dimension embedding 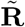 has two advantages. First, it allows us to identify sets of cells with similar copy number profiles, without the overhead of copy number calling. Second, distance functions behave counter-intuitively in high dimensions [46, 47] and therefore dimensionality reduction provides us with a useful proxy for the distance between the copy number profile of pairs of cells. We model the generation of the binned read counts embedding 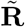 using a Gaussian mixture model where 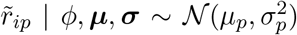. Latent variables 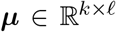 and 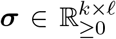 are independent for each clone *j* ∈ [*k*] and dimension *p* ∈ [*ℓ*].

Following convention, let *z_i_* ∈ [*k*] be a random variable depicting the assignment of cell *i* to one of the *k* clones. Given a cell clustering *ϕ*, we have *P*(*z_i_* = *j* | *ϕ*) = **1**{*i* ∈ *ϕ*(*j*)} meaning ϕ specifies a hard clustering of cells into *k* clones. Given cell clustering *ϕ* and latent variables ***μ**_j_* = [*μ_jP_*] and ***σ**_j_* = [*σ_jp_*] for each clone *j* ∈ [*k*], the probability of binned read count embedding for cell *i* and dimension *p* is thus

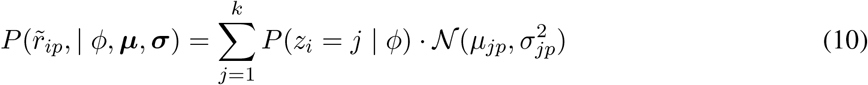

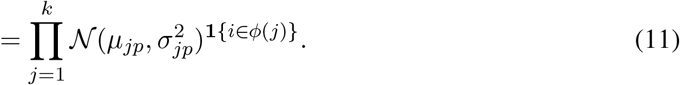

Thus, using independence among cells and dimensions, we have the following probability

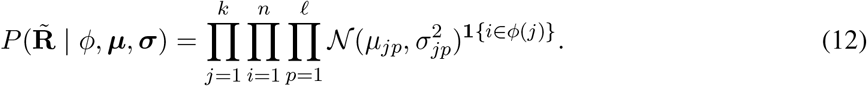

#### A.2.4 Posterior probability

As discussed in Main Text, given hyperparameters *θ* = (*c, α*) we seek to solve

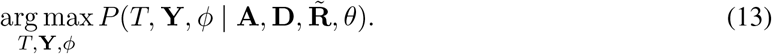

This is equivalent to marginalizing over latent variables ***μ*** and ***σ***,

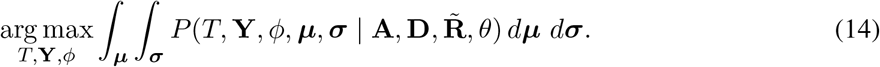

Computing the integrals over latent variables ***μ*** and ***σ*** is challenging in practice, so we instead approximate the above expression by searching for the maximum *a posteriori* (MAP) estimates for latent variables (***μ**, **σ***), giving the expression

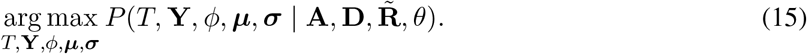

We begin by applying Bayes’ rule, obtaining

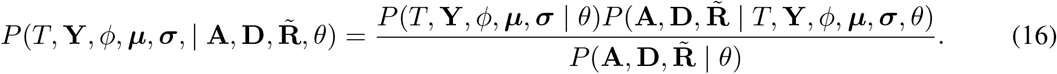

We decompose *P*(*T*, **Y**, *ϕ*, ***μ**, **σ*** | *θ*) per the graphical model and obtain

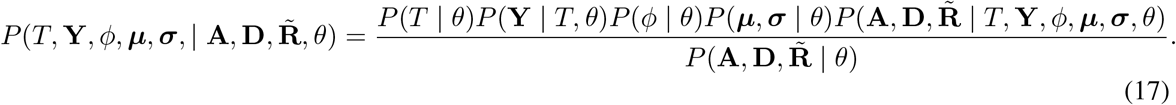

Using uniformity of *P*(*T* | *θ*), *P*(**Y** | *T, θ*), *P*(*ϕ* | *θ*), *P*(***μ**, **σ*** | *θ*) and dropping 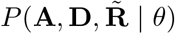 from the denominator, we obtain

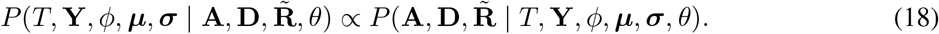

Using the law of conditional probability, we get

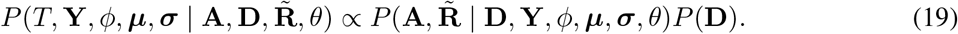

We have that *P*(**D**) is uniform, and after simplification, yields

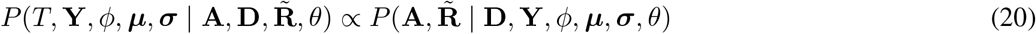

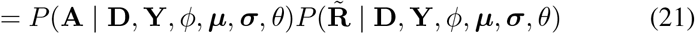

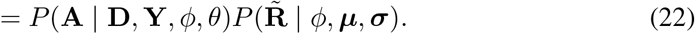

We now focus on *P*(**A** | **D**, **Y**, *ϕ, θ*), which equals

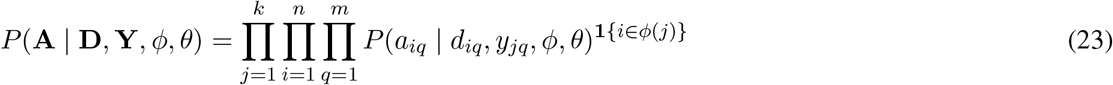

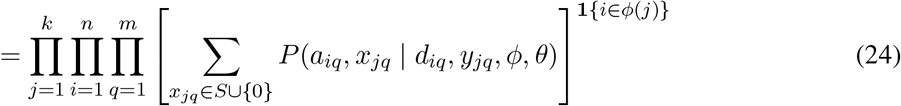

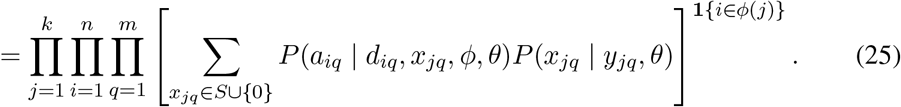

Bringing everything together, we get

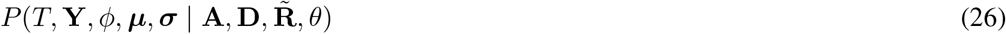

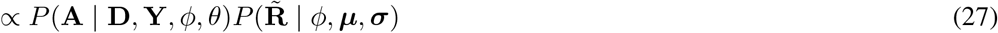

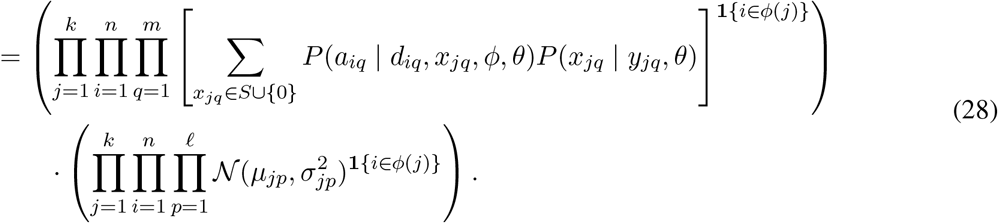

We now take the logarithm, yielding

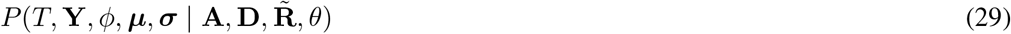

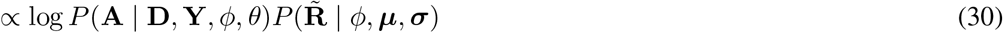

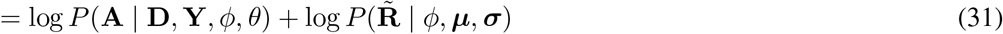

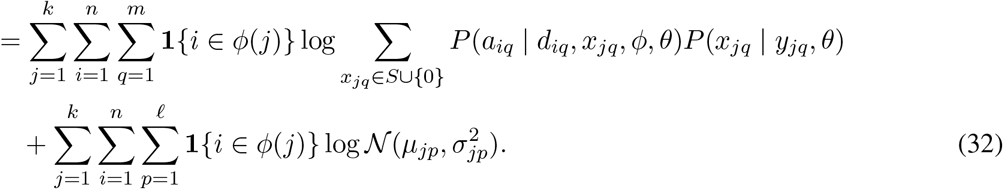

Via the marginalization of latent VAFs **X** and taking the MAP estimates of latent variables ***μ*** and ***σ***, the above expression allows us to approximate the posterior probability 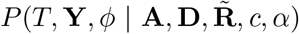 of a given clonal tree T, clonal genotypes **Y** and cell clustering *ϕ*.

### A.3 Detectability of a clone

We define *observations O*(*N, M*) for a subset *N* ⊆ [*n*] of cells and a subset *M* ⊆ [*m*] of SNVs as

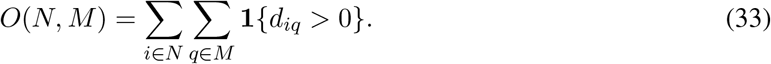

Thus, *O*(*i, M*) is the number of SNV loci in *M* that have mapped reads in cell *i*. Similarly, *O*(*N, q*) is the number of cells in *N* that have mapped reads for SNV locus *q*. Given a threshold parameter 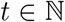, we use this function to define when a clone comprised of cells *N* and newly introduced SNVs *M* is detectable in the following way.

#### Definition 4.

Given parameter 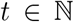, a clone comprised of cells *N* and newly introduced SNVs *M* is *detectable* provided (i) median {*O*(*i, M*) | *i* ∈ *N*} ≥ *t* and (ii) median {*O*(*N, q*) | *q* ∈ *M*} ≥ *t*.

We refer to Fig. S2 for an example.

**Figure S2:**
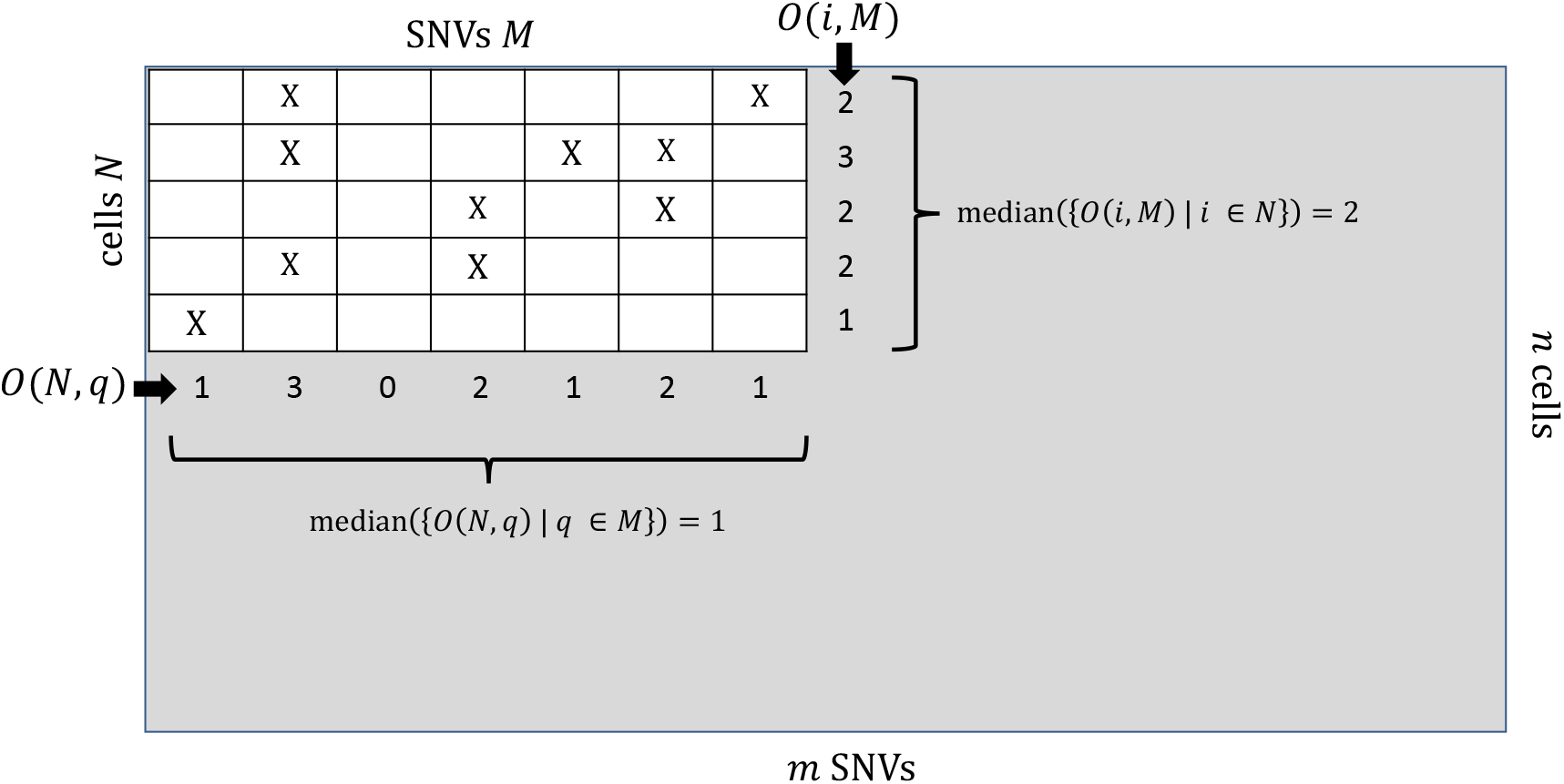
Visual depiction of the detectability of a clone with subset *N* ⊆ [*n*] of cells and subset *M* ⊆ [*m*] of SNVs. Here, ‘X’ indicates the presence of a mapped read for SNV locus *q* in cell *i*. Observations *O*(*N, q*) is the marginal sum of observations over cells *N* for each SNV *q* ∈ *M*. Similarly, *O*(*i, M*) is the marginal sum of observations over SNVs *M* for each cell *i* ∈ *N*. For a clone to be detectable, the medians of these marginal sums must be at least a specified threshold 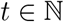 (Definition 4).

### A.4 Phertilizer

At the heart of Phertilizer is the definition of three elementary tree operations: `Branching, Linear` and `Identity`. The input to each operation is a clonal tree *T* with genotypes **Y**, cell clustering *ϕ* and a leaf node *v_j_*. The output is a new clonal tree *T′* with new genotypes **Y′** and cell clustering *ϕ′*. To define these elementary tree operations more formally, we introduce some notations. Given node *v_j_* not equal to the root *v*_1_, the function par(*j*) returns its parent node *v*_par_(*j*) in clonal tree *T*. Recall that y_*j*_ ∈ {0, 1}^*m*^ is the clonal genotype of clone *j* corresponding to node *v_j_* in *T*. With slight overloading of notation, y_*j*_ also indicates the set of SNVs present in clone *j*. We denote the set of SNVs introduced on the incoming edge (*v*_par_(*j*), *v_j_*) of node *v_j_* by Δ(y_*j*_) = y_*j*_ \ y_par_(*j*).

A `Linear` operation transforms node *v_j_* of clonal tree *T* with cells *ϕ*(*j*) and mutations Δ(y_*j*_) into a clonal tree *T′* with a new node *v_a_* and directed edge (*v_j_, v_a_*), partitioning original cells *ϕ*(*j*) and mutations Δ(y_*j*_) among nodes *v_j_* and *v_a_* such that the resulting clonal genotypes 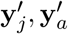 adhere to the infinite sites model, i.e., 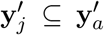 and 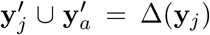 (Main Text Fig. 2b). Similarly, a `Branching` operation results in a clonal tree *T′* with two new nodes *v_a_, v_b_* that are children of *v_j_* such that the original cells *ϕ*(*j*) are partitioned between child nodes *v_a_*, *v_b_*, the mutations Δ(y_*j*_) are partitioned between all three nodes *v_j_, v_a_, v_b_*, and the genotypes 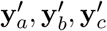 also adhere to the infinite sites model, i.e., 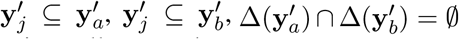 and 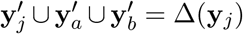. We note that this operation does not assign cells to *v_j_*, thus modeling an extinct clone with SNVs that are common to its children but absent from its parent (Main Text Fig. 2c). Lastly, an `Identity` operation on *v_j_* does not alter the tree and prevents the application of other elementary operations on that node (Main Text Fig. 2d).

Performing an elementary operation on a clonal tree *T* with leaf node *v_j_* is equivalent to solving a constrained CTI problem. That is, we seek to infer a subtree rooted at node *v_j_* with maximum posterior probability that is constrained to either be a two node linear subtree or a three node branching subtree. Both operations use coordinate descent to alternately find a partition of cells and SNVs that maximizes the posterior probability of the inferred clonal subtree. For ease of exposition, we describe the input to these operations as a set *N* = *ϕ*(*j*) of cells assigned to clone *j* in tree *T* and a set *M* = Δ(*v_j_*) of SNVs gained on the incoming edge to node *v_j_*. Next, we describe (i) the `Linear` elementary operation (Appendix A.4.1) (ii) the `Branching` elementary operation (Appendix A.4.2), (iii) a number of regularization steps to avoid overfitting the data (Appendix A.4.3) and (iv) a post-processing phase to improve clonal genotyping and cell clustering in the inferred clonal tree (Appendix A.4.4).

#### A.4.1 Linear

**Figure S3:**
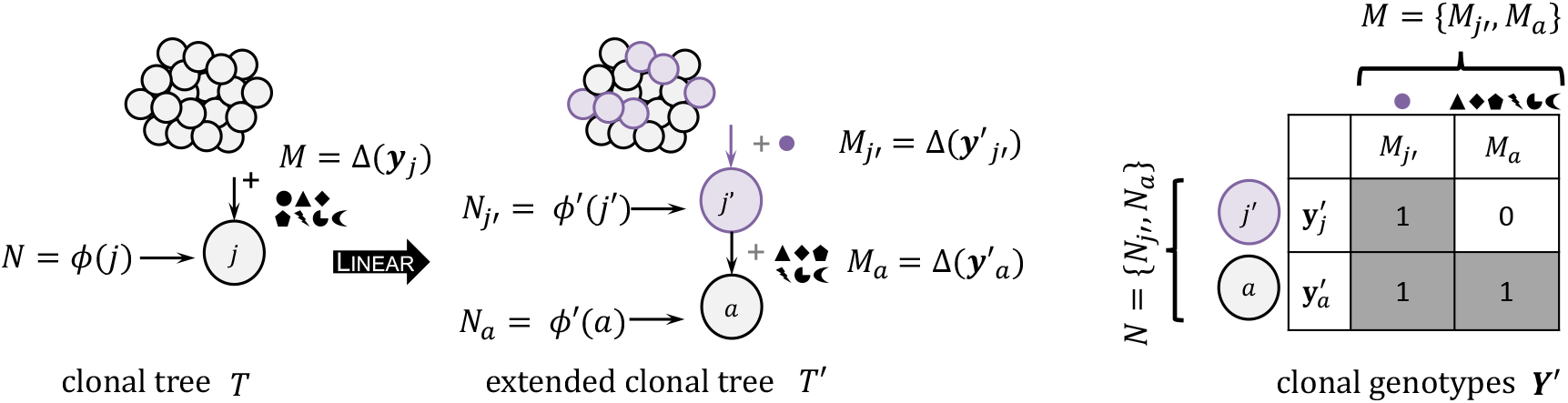
A graphical depiction of a `Linear` elementary tree operation. The cells *N* = *ϕ*(*j*) and the SNVs *M* = Δ(y_*j*_) associated with clone *j* of tree *T* are partitioned into *N* = {*N_j′_*, *N_a_*} and *M* = {*M_j′_*, *M_a_*}, respectively. These partitions are then used to update clonal genotypes **Y′** and cell clustering *ϕ′* associated with extended tree *T′*.

For a given clonal tree *T*, we specify leaf node *v_j_* as the node on which the `Linear` operation is to be performed. The `Linear` operation (Fig. S3) extends tree *T* to create tree *T′* by replacing node *v_j_* of tree *T* by a subtree rooted at node *v_j_* with a single child *v_a_*. Briefly, the `Linear` operation uses a coordinate descent approach to alternately optimize a two-part cell partition and a two-part SNV partition. To optimize the cell partition for a fixed SNV partition, we use the normalized cut algorithm [32]. To optimize the SNV partition for a fixed cell partition, we use our generative model to assign each SNV to the part that maximizes the posterior probability of the extended tree. Next, we describe these steps more formally.

Let *N* = *ϕ*(*j*) ⊆ [*n*] be the cells currently assigned to clone *j* associated with node *v_j_* and let *M* = Δ(y_*j*_) ⊆ [*m*] be the set of SNVs introduced on the incoming edge to node *v_j_* of tree *T*. The output of a `Linear` operation is a two-part partition of the cells *N* into {*N_j′_*, *N_a_*} and a two-part partition of the SNVs *M* into {*M_j′_*, *M_a_*} (Fig. S3). Parts *N_j′_* are the cells associated with node *v_j_* in extended tree *T′* and *N_a_* are the cells associated with the newly added child clone of node *v_j_* in extended tree *T′*. The part *M_j′_* contains SNVs that are introduced on the incoming edge to node *v_j_* in extended tree *T′* and the part *M_a_* correspond to the SNVs newly introduced on the the incoming edge of the child node of *v_j_* after the completion of the operation.

First, we initialize a `Linear` operation by partitioning uniformly at random the set *M* of SNVs into two parts {*M_j′_*, *M_a_*}. Gven the SNV partition {*M_j′_*, *M_a_*}, we use the normalized cut algorithm [32] to find a two part cell partition {*N_j′_*, *N_a_*} of cells *N*. The input to the normalized cut algorithm is a weighed adjacency matrix **W**, where *w_ii′_* relates the similarity of two cells *i* and *i′* in the input set. We set the weight *w_ii′_* to incorporate both an SNV feature f and a CNA feature represented by the binned read count embedding 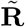.

The SNV feature *f_i_* is defined as 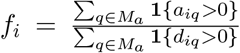. Intuitively, *f_i_* measures for each cell *i* the proportion of introduced SNV loci *M_a_* with mapped reads for which we observe at least one variant read. In case the denominator is 0 for a cell *i*, we set *f_i_* := *f_i″_* where *i″* is the closest cell to *i* in 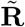 with a nonzero denominator (using Euclidean distance). Then the weight *w_ii′_* between cell *i* and *i′* is defind as

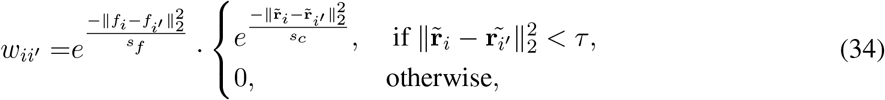

where *τ* is a user-specified parameter. We find that in practice, setting *τ* to the maximum 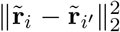 over all pairs of cells *i, i′* ⊆ *N* works well. However, setting *τ* more aggressively, i.e., lower, would put greater emphasis on the CNA features and result in a sparser input to the normalized cut algorithm. Following Shi and Malik [32], parameters *s_f_* and *s_c_* are set to 20% of the total range of the **L**^2^ norm for SNV and CNA features, respectively. We use the `scikit-learn` [48] spectral clustering implementation as it solves the normalized cut problem when the number of clusters is 2. We choose the `cluster-qr` [49] strategy to partition cells in the spectral embedding space.

Then, given the updated cell partition {*N_j′_*, *N_a_*}, we optimize the SNV partition {*M_j′_*, *M_a_*}. Let 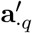 and 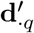 be the column vector of variant and total read counts, respectively, corresponding to cells *N_j′_*. For each SNV *q* ⊆ *M*, we compute the log-likelihood log 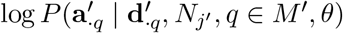 where *M′* ⊆ {*M_j′_*, *M_a_*}. These probabilities are computed as follows.

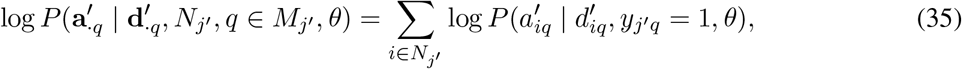

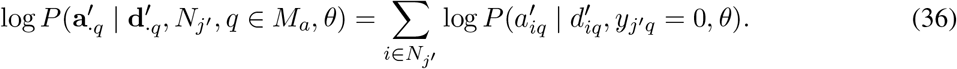

In other words, we compare the probabilities of assigning SNV *q* to either *M_j′_* or *M_a_*. We assign *q* to the part *M′* ⊆ {*M_j′_*, *M_a_*} with maximum log-likelihood. We then iterate until either the cell clusters are not altered from the previous iteration or a user-specified maximum number of iterations is reached (default 50 iterations).

Due to the random initialization of the SNV partition, we repeat this process for a number of restarts. After a restart terminates, we compute a normalized log likelihood term for only the clonal genotypes that will not change even if the leaf nodes of the output tree *T′* are later extended. Among all restarts, the `Linear` operation stores the extended tree *T′*, clonal genotypes **Y′** and cell clustering *ϕ′* with maximum normalized log likelihood. For our simulation study, we found that 16 restarts led to stable results. For experimental data, we conservatively used 25 restarts.

Next, we recurse on an input with cells *N* := *N_j′_* and SNVs *M* ≔ *M_j′_*. The recursion terminates when only an `Identity` operation can be performed. See Appendix A.4.3 for details on this criteria. The recursion step is necessary because the normalized cut algorithm is a recursive partitioning algorithm. Unlike `Branching` (Appendix A.4.2), where all the partitioned cells are associated with leaf nodes in the extended tree nodes, the `Linear` updates the cell cluster of an internal node in the extended tree *T′*. Since these cells will never be taken as input in any further elementary tree operation, recursions helps to refine the clones in the extended tree *T′* as much as possible. After the recursion terminates, the `Linear` operation returns the extended tree *T′*, clonal genotypes **Y′** and cell clustering *ϕ′* with maximum normalized log-likelihood.

#### A.4.2 Branching

**Figure S4:**
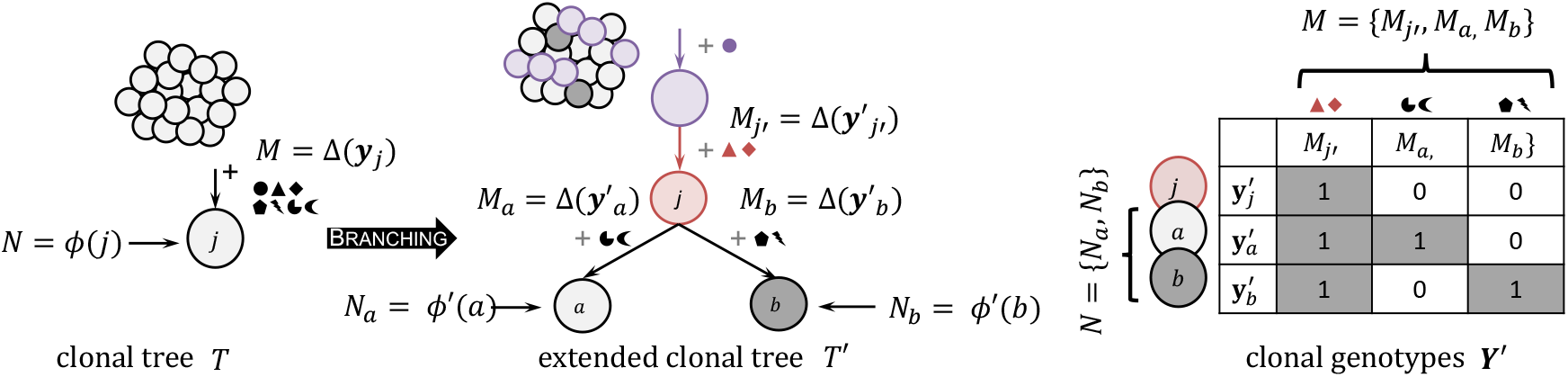
A graphical depiction of a `Branching` elementary tree operation. The cells *N* = *ϕ*(*j*) and the SNVs *M* = Δ(y_*j*_) associated with clone *j* of tree *T* are partitioned into *N* = {*N_a_, N_b_*} and *M* = {*M_j′_*, *M_a_*, *M_b_*}, respectively. These partitions are then used to update clonal genotypes **Y′** and cell clustering *ϕ′* associated with extended tree *T′*.

For a given clonal tree *T*, we specify leaf node *v_j_* as the node on which the `Branching` operation is to be performed. The `Branching` operation (Fig. S4) extends tree *T* to create tree *T′* by replacing node *v_j_* of tree *T* by a binary subtree rooted at node *v_j_* with two children *v_a_* and *v_b_*. Briefly, the `Branching` operation uses a coordinate descent approach to alternately optimize a two-part cell partition and a three-part SNV partition. To optimize the cell partition for a fixed SNV partition, we use the normalized cut algorithm [32]. To optimize the SNV partition for a fixed cell partition, we use our generative model to assign each SNV to the part that maximizes the posterior probability of the extended tree. Next, we describe these steps more formally.

Let *N* = *ϕ*(*j*) ⊆ [*n*] be the cells currently assigned to clone *j* associated with node *v_j_* and let *M* = Δ(y_*j*_) ⊆ [*m*] be the set of SNVs introduced on the incoming edge to node *v_j_* for tree *T*. The output of a `Branching` operation is a two-part partition of the cells *N* into {*N_a_, N_b_*} and a partition of SNVs *M* into three parts {*M_j′_, M_a_, M_b_*} (Fig. S4). Parts *N_a_* and *N_b_* are the cells assigned to the two clones associated with the newly added two children of node *v_j_* in extended tree *T′*. The part *M_j′_* contains SNVs that are introduced on the incoming edge to node *v_j_* in extended tree *T′*. Parts *M_a_* and *M_b_* correspond to the SNVs newly introduced on the two incoming edges of the two children of node *v_j_* after the completion of the operation.

First, we initialize a `Branching` operation by partitioning uniformly at random the set *M* of SNVs into three parts {*M_j′_*, *M_a_*, *M_b_*}. Gven the SNV partition {*M_j′_, M_a_, M_b_*}, we use the normalized cut algorithm [32] to find a two part cell partition {*N_a_, N_b_*} of cells *N*. The input to the normalized cut algorithm is a weighed adjacency matrix **W**, where *w_ii′_* relates the similarity of two cells *i* and *i′* in the input set. We set the weight *w_ii′_* to incorporate both an SNV feature f and a CNA feature represented by the binned read count embedding 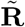. The SNV feature is set as f_*i*_ = [*f_ia_*, *f_ib_*]^T^, where

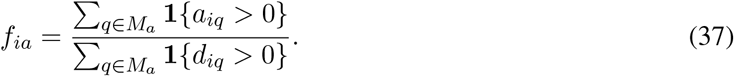

In words, *f_ia_* measures for each cell *i* the proportion of SNVs in set *M_ℓ_* with mapped reads for which we observe at least one variant read. In case the denominator is 0 for a cell *i*, we set *f_ia_* := *f_i″a_* where *i″* is the closest cell to *i* in 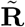 with a nonzero denominator (using Euclidean distance). SNV feature *f_ib_* is similarly defined for SNVs *M_b_*.

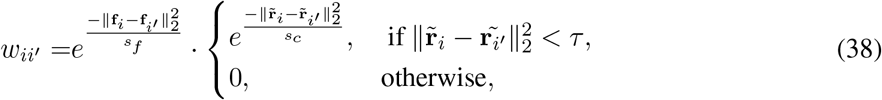

where *τ* is a user-specified parameter. We find that in practice, setting *τ* to the maximum 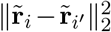 over all pairs of cells *i*, *i′* ∈ *N* works well. However, setting *τ* more aggressively, i.e., lower, would put greater emphasis on the CNA features and result in a sparser input to the normalized cut algorithm. Following Shi and Malik [32], parameters *s_f_* and *s_c_* are set to 20% of the total range of the **L**^2^ norm for SNV and CNA features, respectively. We use the `scikit-learn` [48] spectral clustering implementation as it solves the normalized cut problem when the number of clusters is 2. We choose the `cluster-qr` [49] strategy to partition cells in the spectral embedding space.

Then, given the updated cell partition {*N_a_, N_b_*}, we optimize the SNV partition {*M_j′_*, *M_a_*, *M_b_*}. Let 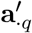 and 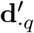 be the column vector of variant and total read counts, respectively, corresponding to cells *N*. For each SNV *q* ∈ *M*, we compute the log-likelihood log 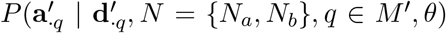 where *M′* ∈ {*M_j′_, M_a_, M_b_*}. These probabilities are computed as follows.

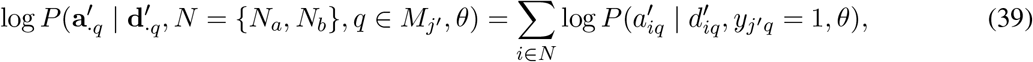

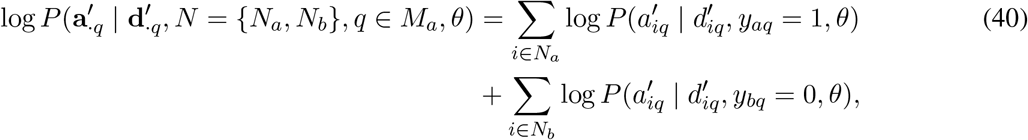

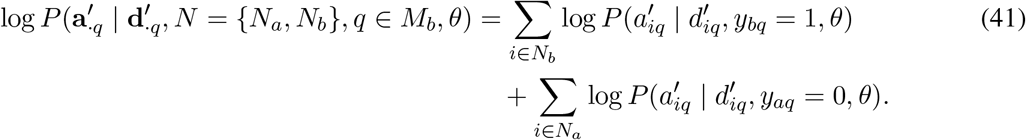

In other words, we compare the probabilities of assigning SNV *q* to either *M_j′_*, *M_a_* or *M_a_*. We assign *q* to the part *M′* ∈ {*M_j′_, M_a_, M_b_*} with maximum log-likelihood. We then iterate until either the cell clusters are not altered from the previous iteration or a user-specified maximum number of iterations is reached.

Due to the random initialization of the SNV partition, we repeat this process for a number of restarts. After a restart terminates, we compute a normalized log likelihood term for only the clonal genotypes that will not change even if the leaf nodes of the output tree *T′* are later extended. Among all restarts, the `Branching` operation returns the extended tree *T′*, clonal genotypes **Y′** and cell clustering *ϕ′* with maximum normalized log likelihood. For our simulation study, we found that 16 restarts led to stable results. For experimental data, we conservatively used 25 restarts.

#### A.4.3 Regularization

To avoid overfitting the input data, we execute a number of regularization steps after performing an elementary tree operation on a node *v_j_* of clonal tree *T*.

##### Detectability check

We assess whether the clones associated with new nodes *v**j**, v_a_, v_b_* for `Branching` (Appendix A.4.2) and nodes *v_j_, v_a_* for `Linear` (Appendix A.4.1) are detectable (Appendix A.3). We set the default as *t* = 4.

##### Quality assessment of an extension

Even though we may have enough data observations to theoretically perform an elementary operation on a leaf node *v_j_* in tree *T*, a high quality extension might not exist. We define the *cell mutational burden* (CMB) for a cell *i* ∈ *N* ⊆ [*n*] and set *M* ⊆ [*m*] of SNVs as

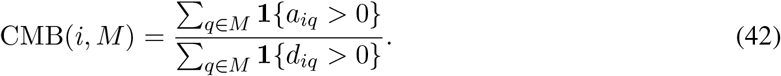

More details and intuition on the CMB is provided in Appendix B.2.3.

We define a quality check qc(*N, M*) as median{CMB(*i, M*) | *i* ∈ [*N*]}. When attempting a `Linear` operation, which partitions *N* = *ϕ*(*j*) into parts {*N_j′_*, *N_a_*} and SNVs *M* = Δ(y_*j*_) into parts {*M_j′_, M_a_*} (Appendix A.4.1) we expect qc(*N, M_j′_*) to be high, i.e., at least 0.15, since we expect all *M_j′_* SNVs to be present in the set *N* of cells. Conversely, we expect qc(*N_j′_*, *M_a_*) to be low, i.e., at most 0.05, since we do not expect to see SNVs *M_a_* in the set *N_j′_* of cells. For similar reasoning as above, we define the following quality checks for the `Branching` operation (Appendix A.4.2).

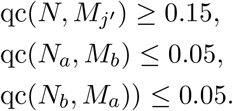

We use 0.05 and 0.15 for illustrative purposes and because we set these as the default values. However, the user may specify different values for these parameters to be less conservative.

#### A.4.4 Postprocessing

Recall that the input to an elementary operation is a node *v_j_* with attached cells *ϕ*(*j*) and introduced SNVs *Δ*(y_*j*_). It may be the case that a cell *i* in *ϕ*(*j*) has zero variant reads for all SNVs in Δ(y_*j*_), i.e., *a_iq_* = 0 for all SNVs *q* ∈ Δ(y_*j*_). Similarly, an SNV *q* in Δ(y_*j*_) may not have any cell in *ϕ*(*j*) with supporting variant reads, i.e., *a_iq_* = 0 for all cells *i* ∈ *ϕ*(*j*). These SNVs and cells are uninformative during the elementary operations and their placement within the tree is best determined after the clonal tree has been identified. As such, prior to performing an elementary operation on node *v_j_*, we remove such SNVs and cells, and add them to a set *M* of SNVs and a set *N* of cells, respectively. After Phertilizer returns the clonal tree with maximum posterior probability, we then perform an optional post-processing phase that contains two stages: (i) the placement of SNVs and cells that were removed from the tree during inference and (ii) the reassignment of SNVs that fit poorly in the tree.

For post-processing, we are given a clonal tree *T*, clonal genotypes **Y** and cell clustering *ϕ*, as well as the cells *N* and SNVs *M* as described above. In the first stage, we first fix the cell clustering *ϕ*. Recall that Δ(y_*j*_) is the set of SNVs introduced on the incoming edge to node *v_j_* in tree *T*. Then, for each SNV *q* ⊆ *M* and each node *v_j_* ⊆ *V* (*T*), we compute the likelihood that SNV *q* is gained at node *v_j_* in tree *T* as follows.

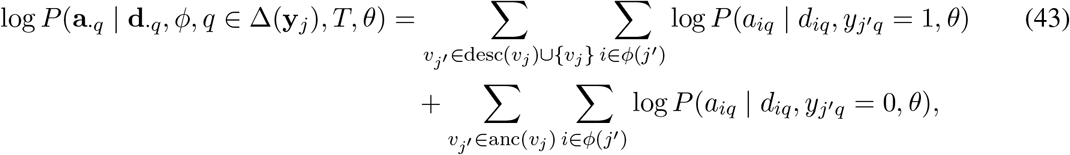

where desc(*v_j_*) is the set of descendant nodes and anc(*v_j_*) is the set of ancestor nodes of node *v_j_* in tree *T*. We assign *q* to the incoming edge of node *v_j_* ⊆ *T* with maximum log-likelihood, resulting in updated clonal genotypes **Y′**. We then set clonal genotypes **Y** ≔ **Y′**.

Next, we repeat the above process but fix the clonal genotypes **Y** and update the cell clustering *ϕ*. For each cell *i* ⊆ *N* and for each node *v_j_* ⊆ *V*(*T*) such that 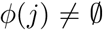, we compute the log-likelihood of cell *i* being assigned to clone *j* as follows.

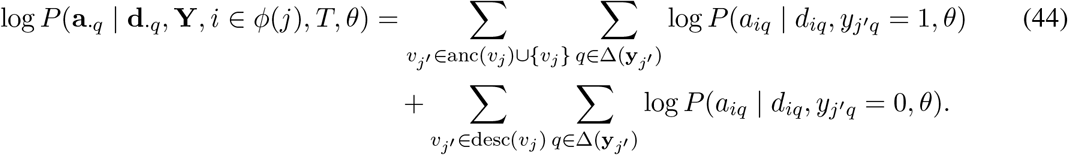

We assign cell *i* to the cell cluster *j* ∈ [*k*], with maximum log-likelihood resulting in updated cell clustering *ϕ′*. We then set cell clusterings *ϕ* ≔ *ϕ′*.

In the second stage, given a tree *T* and a cell clustering *ϕ*, we attempt to improve clonal genotypes **Y** by moving SNVs that fit poorly with the given cell clustering *ϕ* and tree *T*. We iterate over the nodes of tree *T* using a postorder traversal. At each node *v_j_*, we identify poorly fitting SNVs by computing a metric we refer to as the binary variant alele frequency (BVAF), which is as follows for a given SNV *q* and node *v_j_*

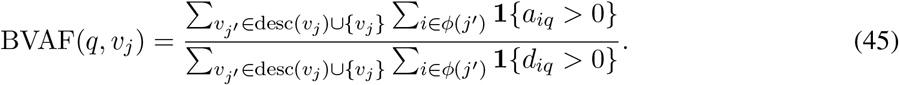

In words, BVAF(*q, v_j_*) is the fraction of cells in clade *v_j_* with mapped variant reads at SNV *q*. We compute BVAF(*q, v_j_*) for every SNV *q* ∈ Δ(y_*j*_). We identify a cutoff *t′* at the 10th percentile of this distribution. Then, any SNV q where BVAF(*q, v_j_*) ≤ *t′* is removed from the tree and placed in a candidate set *M* of SNVs to be reassigned. Reassignment then proceeds as described above.

We repeat this second stage of post-processing for either a fixed number of iterations or until no SNVs are reassigned, resulting in updated clonal genotypes **Y′**. After termination, we update clonal genotypes **Y** ≔ **Y′** and return clonal genotypes **Y** and cell clustering *ϕ*.

## B Supplementary results

### B.1 Simulation study

Below we provide pertinent details related to our simulation study and supplemental figures.

- Appendix B.1.1 provides additional details for the **Baseline** comparison method
- Appendix B.1.2 provides additional details on our simulation study design
- Appendix B.1.3 provides the specification of runtime parameters for methods analyzed in the simulation study
- Appendix B.1.4 provides additional details on the performance metrics used to evaluation the results of our simulation study
- Appendix B.1.5 contains supplemental results figures for our simulation study

#### B.1.1 Baseline method

The Baseline method seeks to infer clonal genotypes for cells utilizing copy-number information embedded in the binned read counts **R** via the following three steps. First, the method projects binned read counts 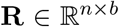 into a low dimensional space 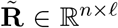 (i.e., *ℓ* ≪ *b*) — see Appendix A.1. Second, it clusters cells into clones in the low-dimensional binned read count embedding 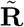. Third, each cluster of cells is treated as a pseudobulk sample and the cluster is genotyped. In our implementation, we used OPTICS [50] for the clustering step, since it is in the family of DBSCAN clustering methods that is a common choice used for the Baseline in practice and the number of clusters does not need to be specified *a priori*. To genotype a cell cluster *j*, we set clonal genotype *y_jq_* = 1 for each SNV *q* ∈ [*m*] whenever the variant allele frequency, i.e., the total number of variant reads/total number of reads, was greater than 0.05 and *y_jq_* = 0 otherwise. Fig. S5 shows good concordance of clonal genotypes using this approach with that of the inferred clonal genotypes on a DLP+ dataset [6].

#### B.1.2 Simulation setup

We generated scDNA-seq data from *in silico* heterogeneous tumors with both CNAs and SNVs. We generated a tree *T*\* with *k* ∈ {5, 9} nodes. We varied the number *n* ∈ {1000, 2000} of sequenced cells and the number m 2 {5000, 10000, 15000} of SNVs. We generated read counts following a beta-binomial distribution with coverage of 0.01×, 0.05×, and 0.1×. We introduced 3 chromosome-level and 8 chromosome-arm level CNAs, affecting a total of *ℓ* = 577 bins each of length 5 MB spanning all autosomes. We sampled a cell clustering *ϕ*\* as well as SNV and CNA placement on edges from a symmetric Dirichlet distribution with a concentration parameter of 2.

Each combination of simulation parameters was replicated with ten different random number generator seeds, amounting to a total of 180 experiments. In addition, we generated a set of simulation experiments in the same manner as above with exception of fixing the copy number as heterzygous diploid at every locus in the genome. We also varied the SNV evolutionary model, using either a Dollo (SNV loss, [34]) or an infinite sites model (without SNV loss, [30]). We used a mean sequencing coverage of 0.01× drawn from a Poisson distribution, to generate data matrices **D**, **R** and sampled the variant count matrix **A** from a binomial distribution parameterized by the ground truth VAF.

#### B.1.3 Runtime parameters

##### Phertilizer

We used default parameters specified below for all simulation instances.

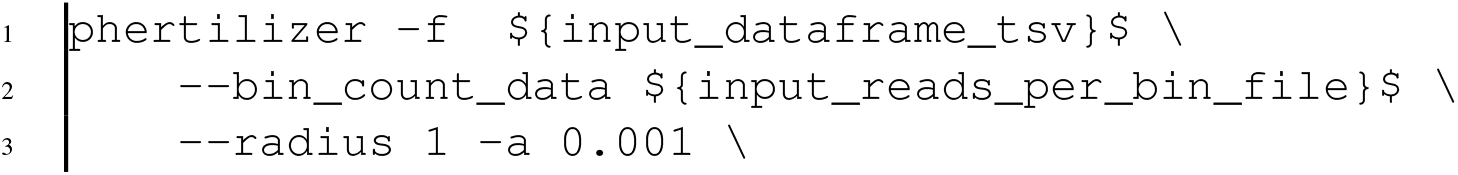

**Figure S5:**
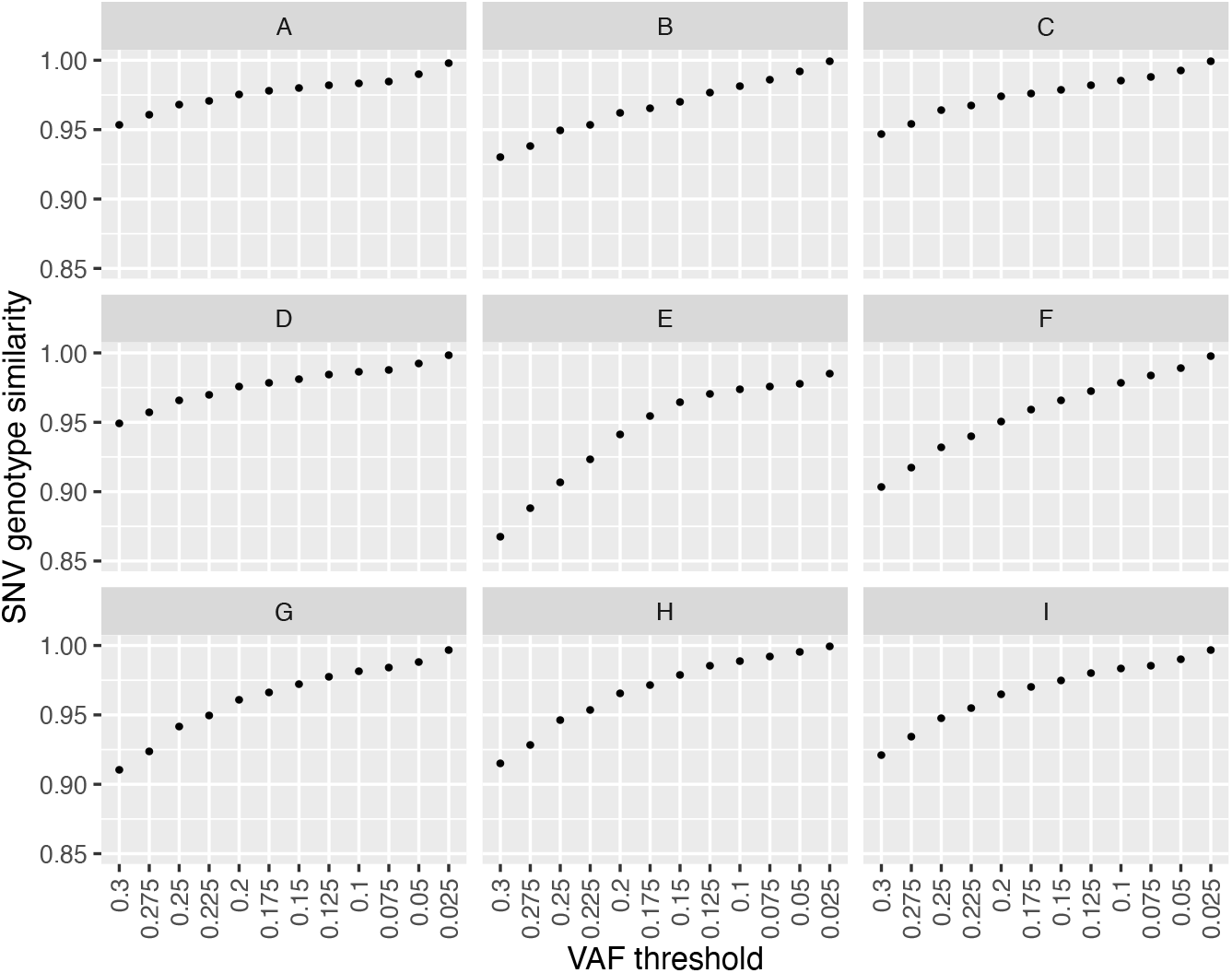
Genotype similarity between inferred genotypes per cell cluster using the **Baseline** method and varying the variant allele frequency (VAF) threshold and inferred clonal genotypes per cell cluster by Laks et al. [6]

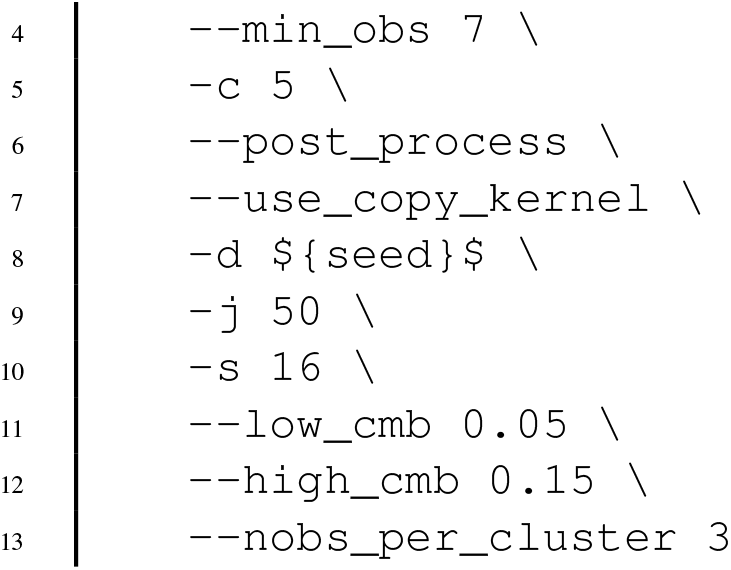

##### SBMClone

We used the same seed for simulation as the seed for SBMClone.

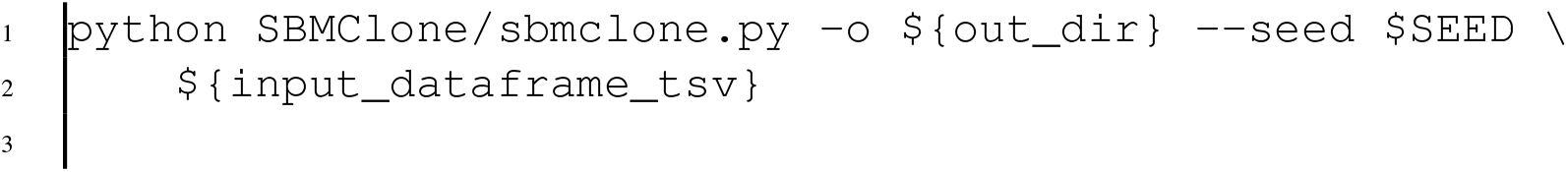

##### SCITE

We set the maximum MCMC chain length to be 900, 000, repetitions to 3, false positive rate to be 0.02, false negative rate to be 0.01, and seed to 42.

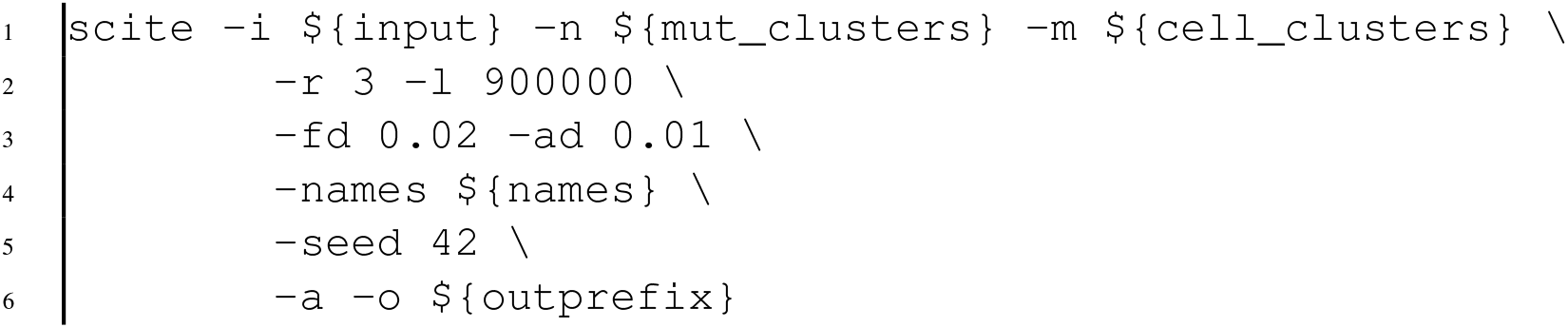

#### B.1.4 Performance metrics

**Figure S6:**
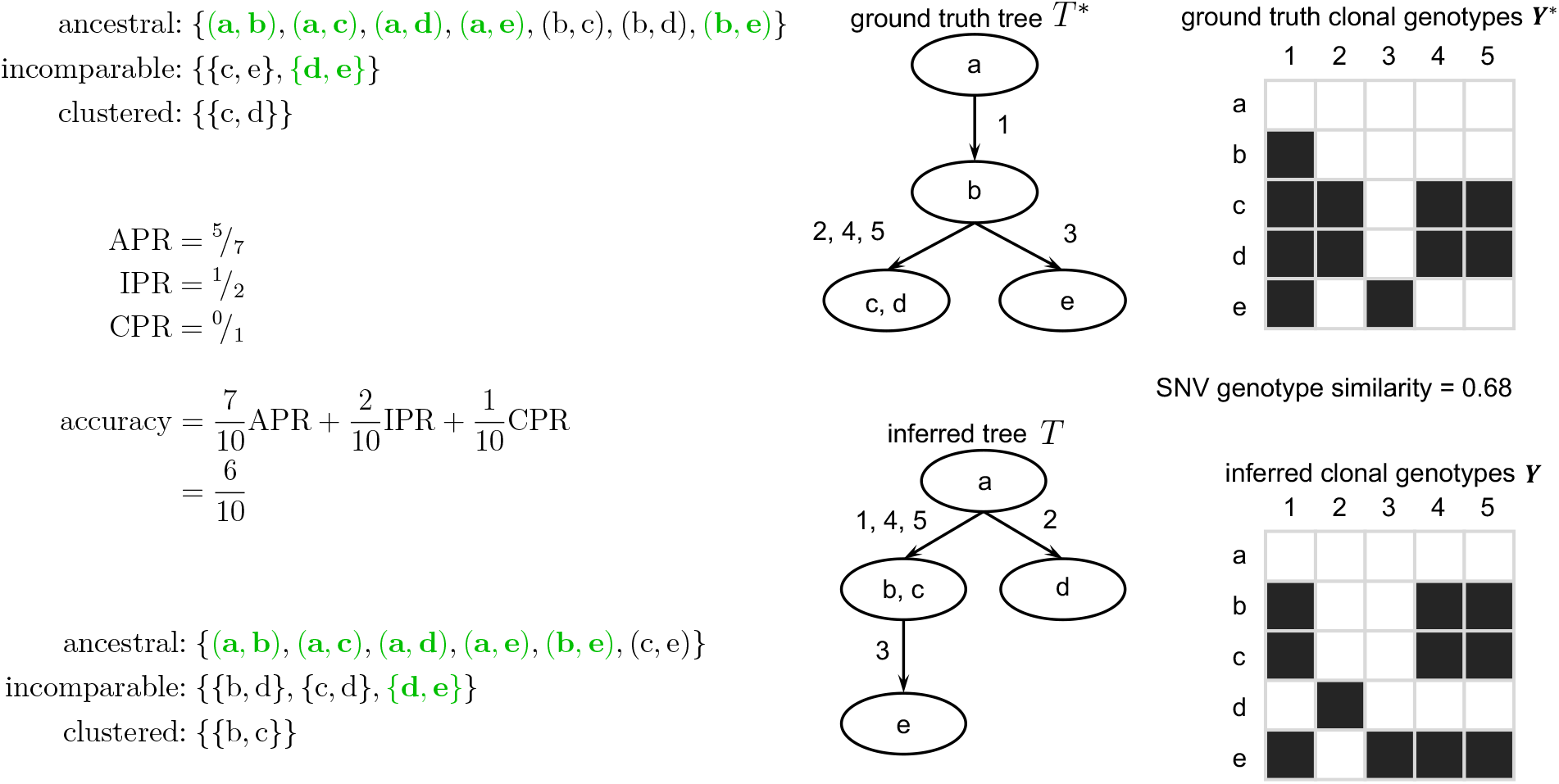
Example for ancestral pair recall (APR), clustered pair recall (CPR), incomparable pair recall (IPR), and accuracy for cells. To the right of the trees, we show the ground truth and inferred genotypes **Y**\* and **Y** projected down to individual cells. Main Text Fig. provides an example of APR, CPR, IPR and accuracy metrics for SNVs.

We assessed the quality of an inferred solution (*T, ϕ*, **Y**) against a ground-truth tree *T*\*, cell clustering *ϕ*\* and clonal genotypes **Y**\* using ancestral pair recall (APR), incomparable pair recall (IPR), and clustered pair recall (CPR) metrics for cells and SNVs [33], as well as genotype similarity. In addition, we compute a single *accuracy* value (∈ [0, 1]) composed of the weighted average of APR, IPR and CPR — where the weights are proportional to the number of pairs in each class. We note that an SNV and cell accuracy of 1 implies that the inferred solution perfectly matches ground truth.

##### APR, IPR, CPR

For any two SNVs *q* = *q′* there are three possible placements in the tree: (i) *ancestral: q* is gained on a node that is distinct and ancestral to the node where *q′* is gained; (ii) *clustered: q* and *q′* are both gained on the same node; and (iii) *incomparable: q* and *q′* are gained on distinct nodes that occur on distinct branches of the tree. Similarly, two distinct cells *i* ≠ *i′* have the same three possible placements. The APR assesses the ratio of ancestral pairs from *T*^*^ recalled in *T*, whereas CPR and IPR do so for clustered and incomparable pairs, respectively. Thus, if APR = CPR = IPR = 1 for both cells and SNVs then the inferred solution (*T, ϕ*, **Y**) is identical to the ground truth (*T*\*, *ϕ*\*, **Y**\*). Main text Fig. 3a and Fig. S6 provide a graphical depiction of these metrics.

##### Genotype similarility

We define the *genotype similarity* as 1 minus the normalized Hamming distance between ground truth genotypes and inferred genotypes of cells. More formally, this is defined as 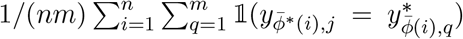 where 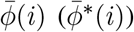 is the unique clone *j* such that *i* ∈ *ϕ*(*j*) (*i* ∈ *ϕ*\*(*j*)), given an inferred genotype **ŷ**_*i*_ ∈ {0, 1}^*m*^ and a ground truth genotype 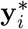. Thus, a genotype similarity of 1 implies that every sequenced cell was correctly genotyped. Fig. S6 above provides a visualization for the computation of this metric.

#### B.1.5 Supplemental simulation study figures

- Fig. S7 shows supplemental simulation study results for coverage 0.01×
- Fig. S8 shows supplemental simulation study results for coverage 0.05×
- Fig. S9 shows supplemental simulation study results for coverage 0.1×
- Fig. S10 shows heterozygous diploid simulation study results
- Fig. S11 shows Dollo evolutionary model simulation study results
- Fig. S12 shows a running time comparison for the simulation study

**Figure S7:**
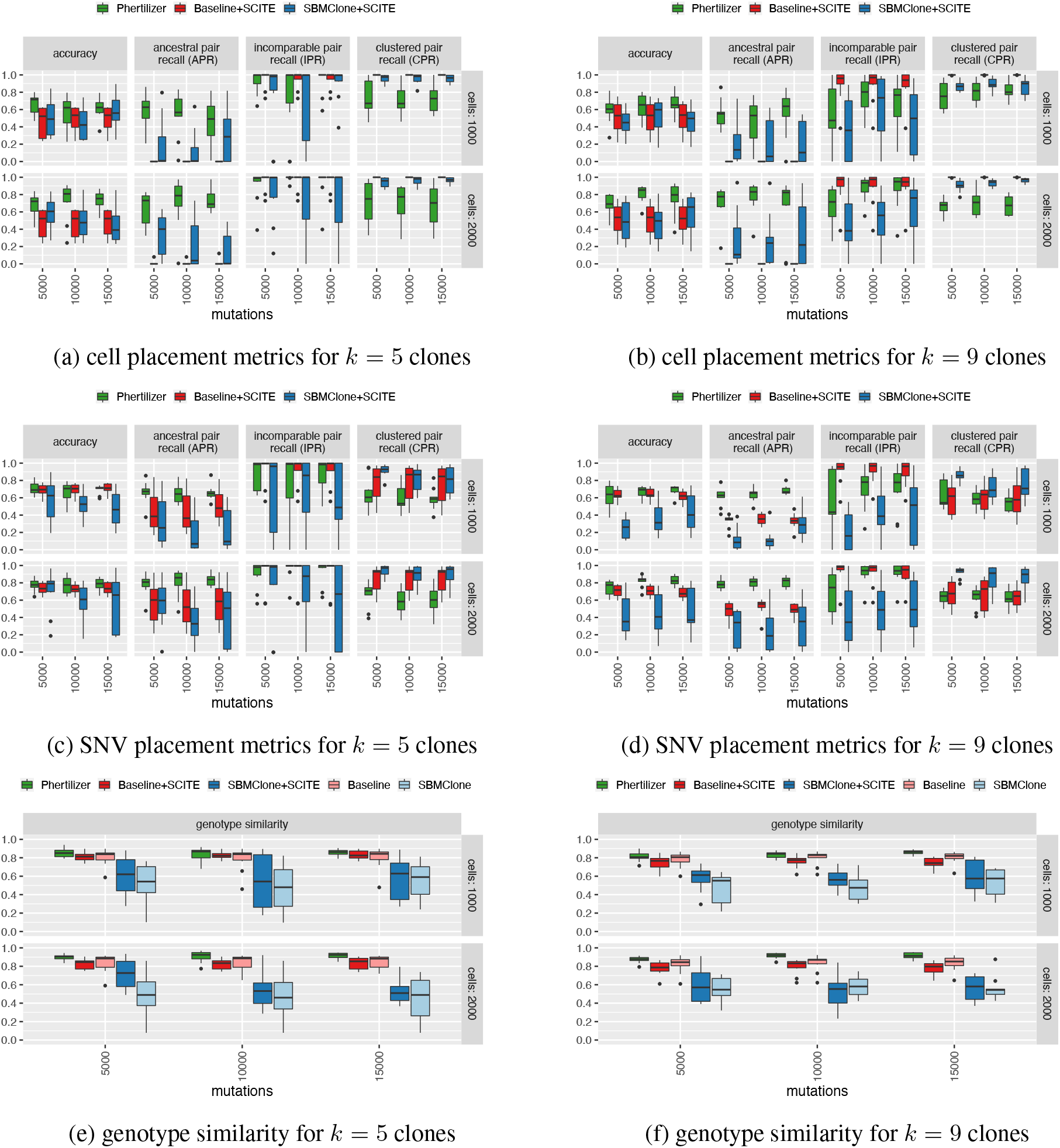
Simulation results for coverage *g* = 0.01 ×

**Figure S8:**
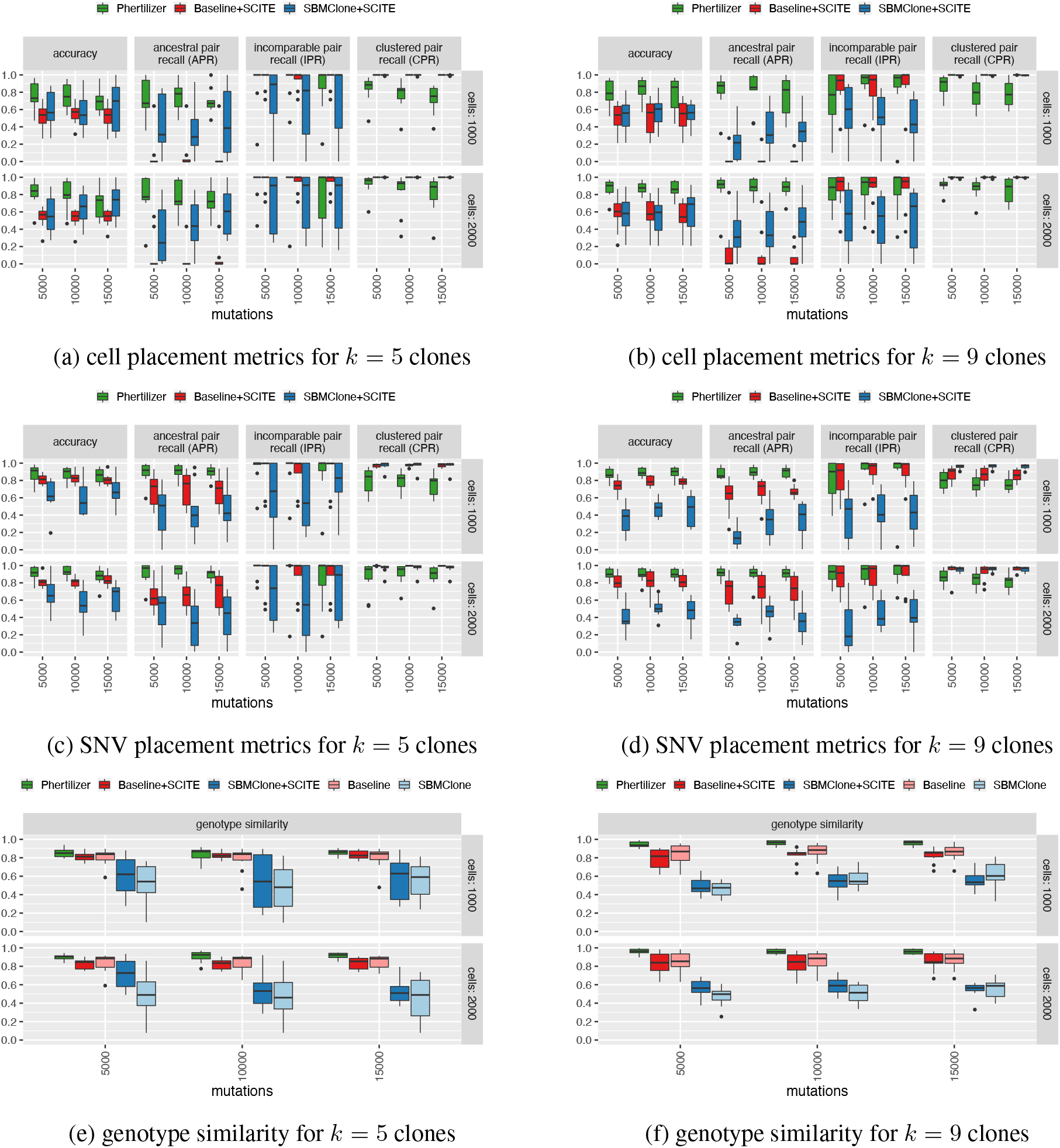
Simulation results for coverage *g* = 0.05×.

**Figure S9:**
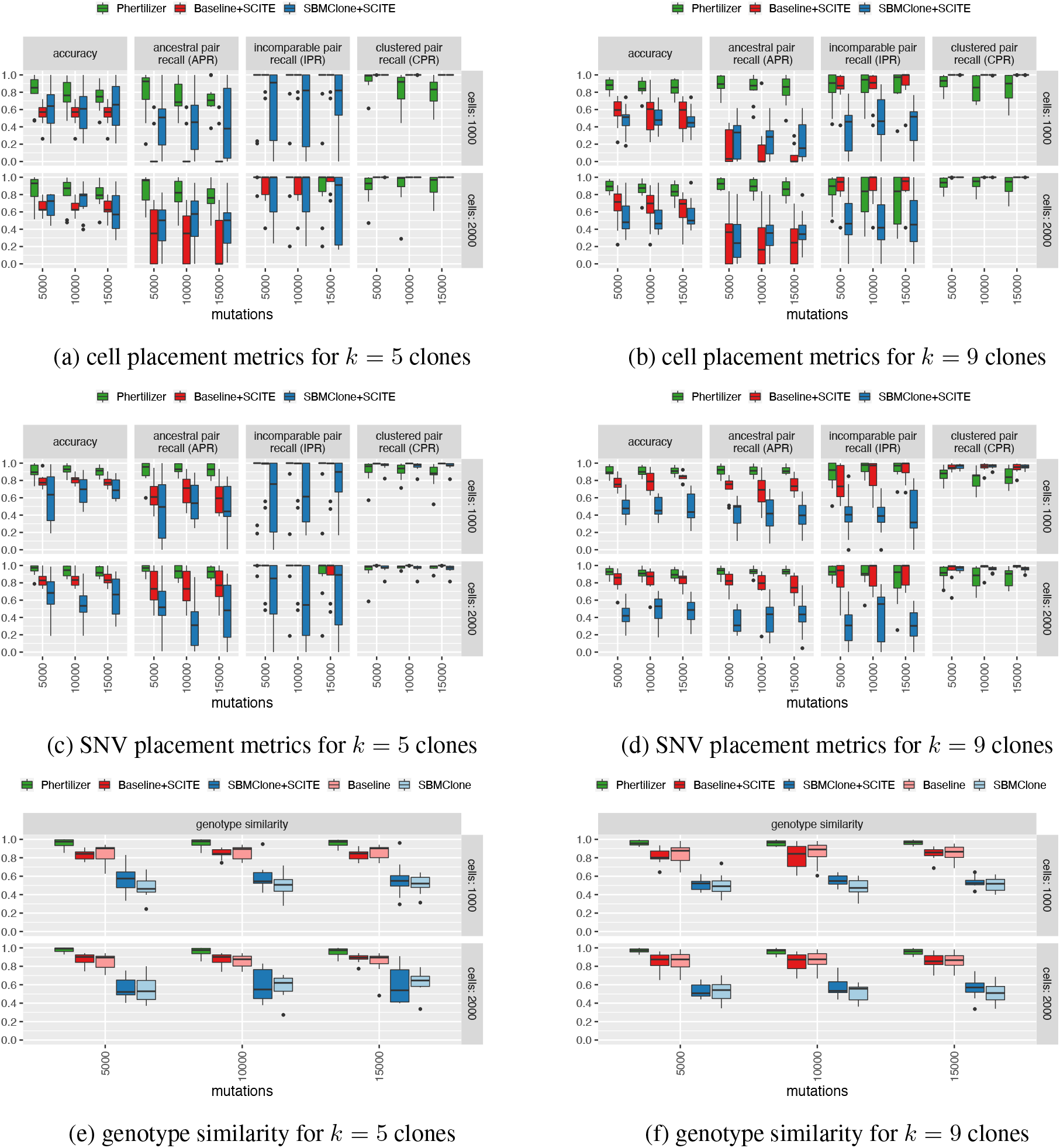
Simulation results for coverage *g* = 0.1×.

**Figure S10:**
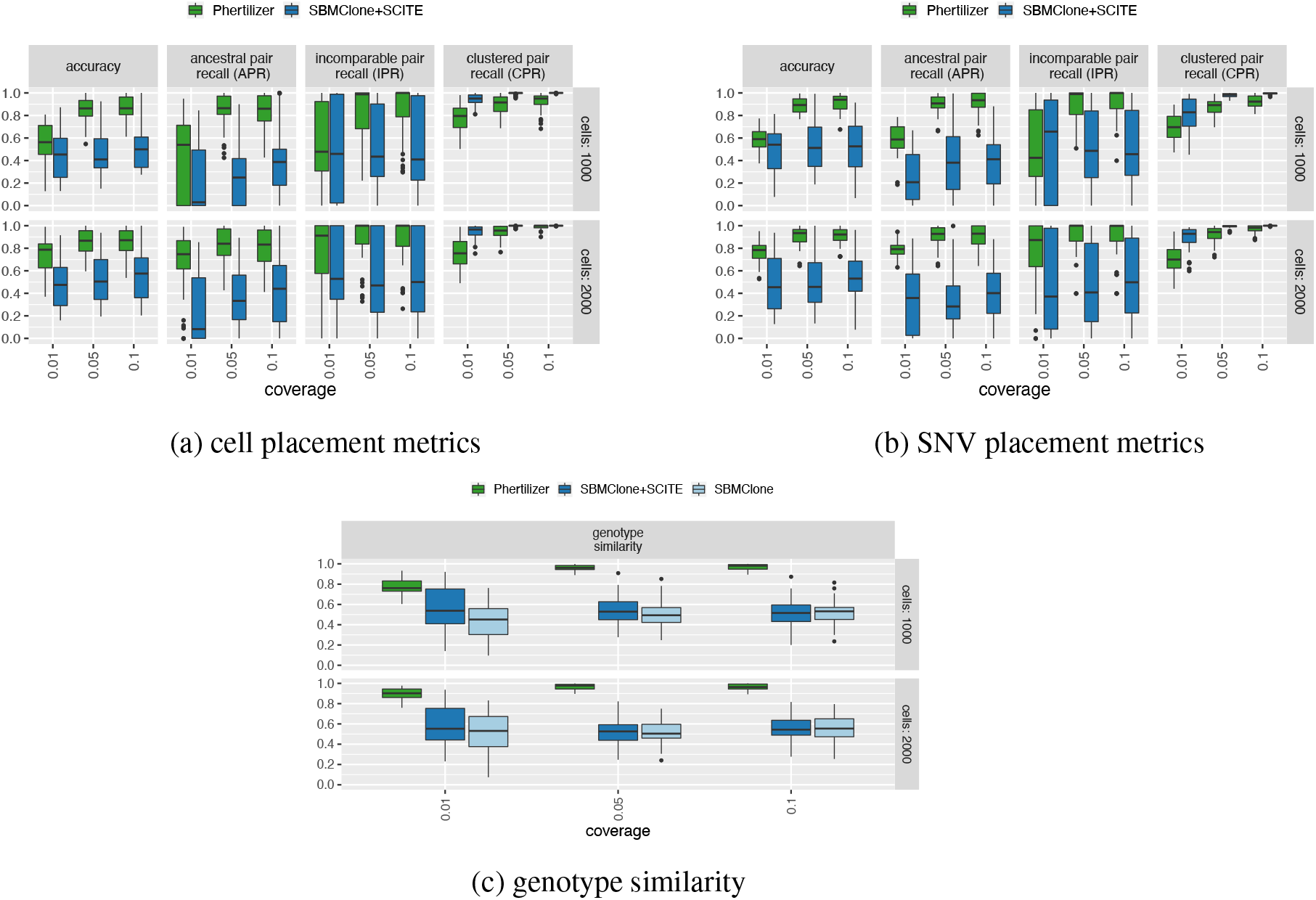
Heterozygous diploid simulation results aggregrated over *k* ∈ {5, 9} clones and *m* ∈ {5000, 10000, 15000} SNVs. Baseline+SCITE was excluded from comparison because the lack of CNAs in the data resulted in only a single clone being identified in the read count embedding space for each instance.

**Figure S11:**
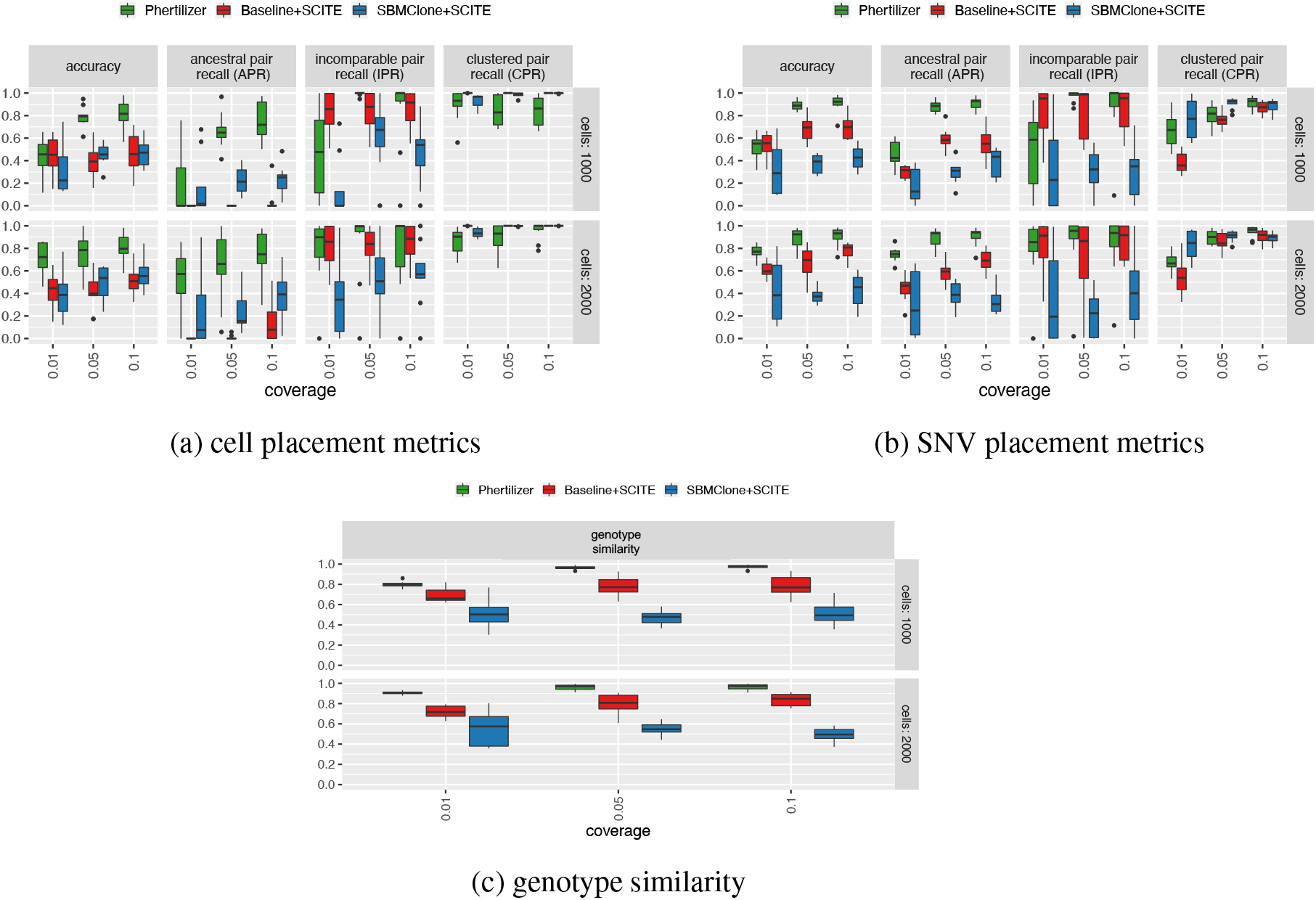
Dollo evolutionary model simulation results for *k* = 9 clones and *m* = 15000 SNVs.

**Figure S12:**
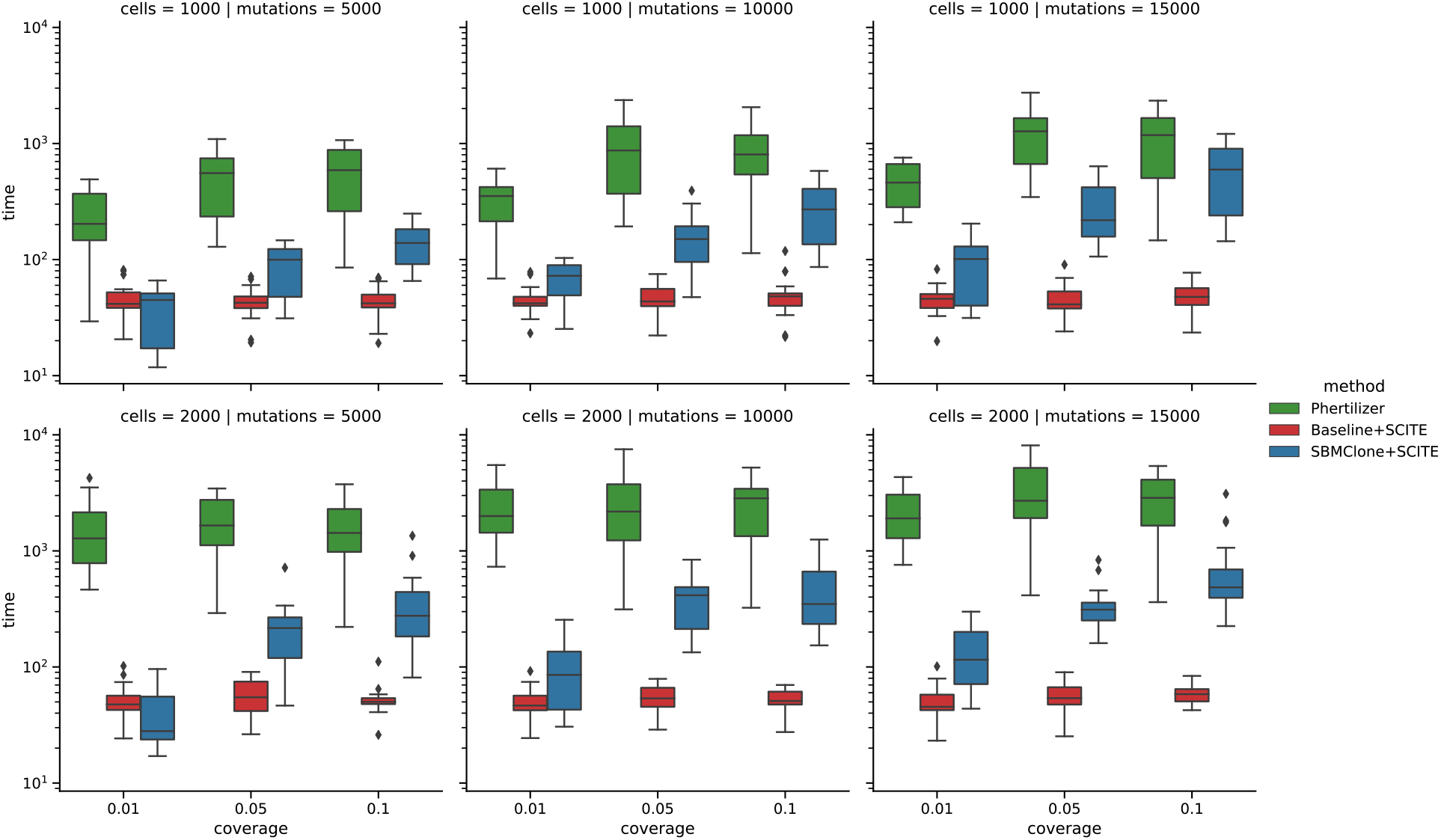
Running time (in seconds) on simulation data.

### B.2 Experimental data

Below we provide pertinent details and supplemental results for our analysis of high-grade serous ovarian cancer cells sequenced via DLP+ and triple negative breast cancer tumors sequenced via ACT.

- Appendix B.2.1 provides details of the Phertilizer runtime parameters used for experimental data.
- Appendix B.2.2 contains amplifying information on how cancer related genes were placed on inferred trees.
- Appendix refsupp: cell mutational burden provides additional description and analysis of the cell mutational burden metric
- Appendix B.2.4 contains supplementary results for the high-grade serous ovarian cancer cells sequenced with DLP+
- Appendix B.2.5 provides the data processing details for the triple negative breast cancer tumors sequenced via ACT
- Appendix B.2.6 contains supplemental results and figures for the triple negative breast cancer tumors sequenced via ACT

#### B.2.1 Phertilizer runtime parameters for experimental data

We set hyperparameters *c* = 5 and *ρ* = 0.001 for all experimental data analyses. We used 25 restarts and a maximum of 50 iterations for each elementary tree operation. Additionally, we performed a grid search over detectability threshold *t* ∈ {5, 6, 7} (Appendix A.3) and the lower bound for the quality check qc ∈ {0.05, 0.075} (Appendix A.4.3). The upper bound for the quality check qc was set to 0.15. After running Phertilizer on each of these 6 combinations of parameters, we selected the clonal tree with maximum posterior probability.

#### B.2.2 Placement of driver genes on inferred trees

We annotated inferred clonal trees with cancer-related genes listed in the Cancer Gene Census (CGC) [37] from COSMIC v97 and cBioPortal [35, 36]. In particular, we identified missense variants in cancer-related genes as well as genes with stop-gain variants. We first selected stop-gain variants. Then, we annotated variants with VEP [51], and looked for missense variants that are predicted to be damaging or deleterious by both SIFT [52] and PolyPhen [53]. Among these, we selected variants either present in the CGC, or in more than 0.4% of the total 76,639 samples from 10 pan-cancer studies included in cBioPortal. As a large number of SNVs are found in patient TN8 of the ACT dataset, we only labeled genes present in the CGC.

#### B.2.3 Cell mutational burden (CMB)

To assess the quality of each inferred clade for real data, we developed a performance metric called *cell mutational burden* (CMB) defined as

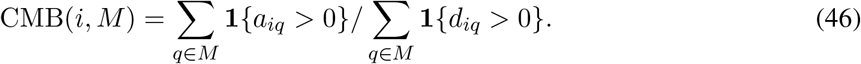

In words, CMB(*i, M*) is the fraction of mapped SNV loci *M* with mapped variant reads in cell *i*. For a specified *clade j* or subtree rooted at node *v_**j**_*, SNVs *M_**j**_* are the SNVs gained at node *v_**j**_*. CMB is designed to assess the goodness of fit of a proposed clonal tree without a known ground truth tree and succinctly captures the relationship between the inferred tree, clonal genotyping, and cell clustering. For a cell *i* placed within clade *j*, we expect CMB(*i, M_**j**_*) to be high, although the value will depend on copy number. By contrast, for cells placed outside of clade *j*, we expect CMB(*i, M_**j**_*) to be low.

Because this is a newly proposed metric that has complex interactions between inference errors, we performed a sensitivity analysis on a simulated tree (Fig. S13a) with varying rates in {0%, 15%, 30%} of cell and SNV placement errors (Fig. S13). For reference, we also show cell and SNV placement on the simulated ground truth tree (Fig. S13b) at the highest error rate for both (30%). This analysis showed that in the error free regime, CMB value for cells in clade ranges from from 0.2 to 0.8 but the value will depend on copy number. We also found that as SNV error rates increase, the median CMB for cells outside of the clade increases with median values as high as 0.1 and an increased number of outliers with values as high as 0.7. Finally, as cell error rates increase the variability of CMB values for cells in clade increases. Although this analysis was only performed on a single simulation instance, it is helpful for interpreting this metric on experimental data where ground truth is unknown.

**Figure S13:**
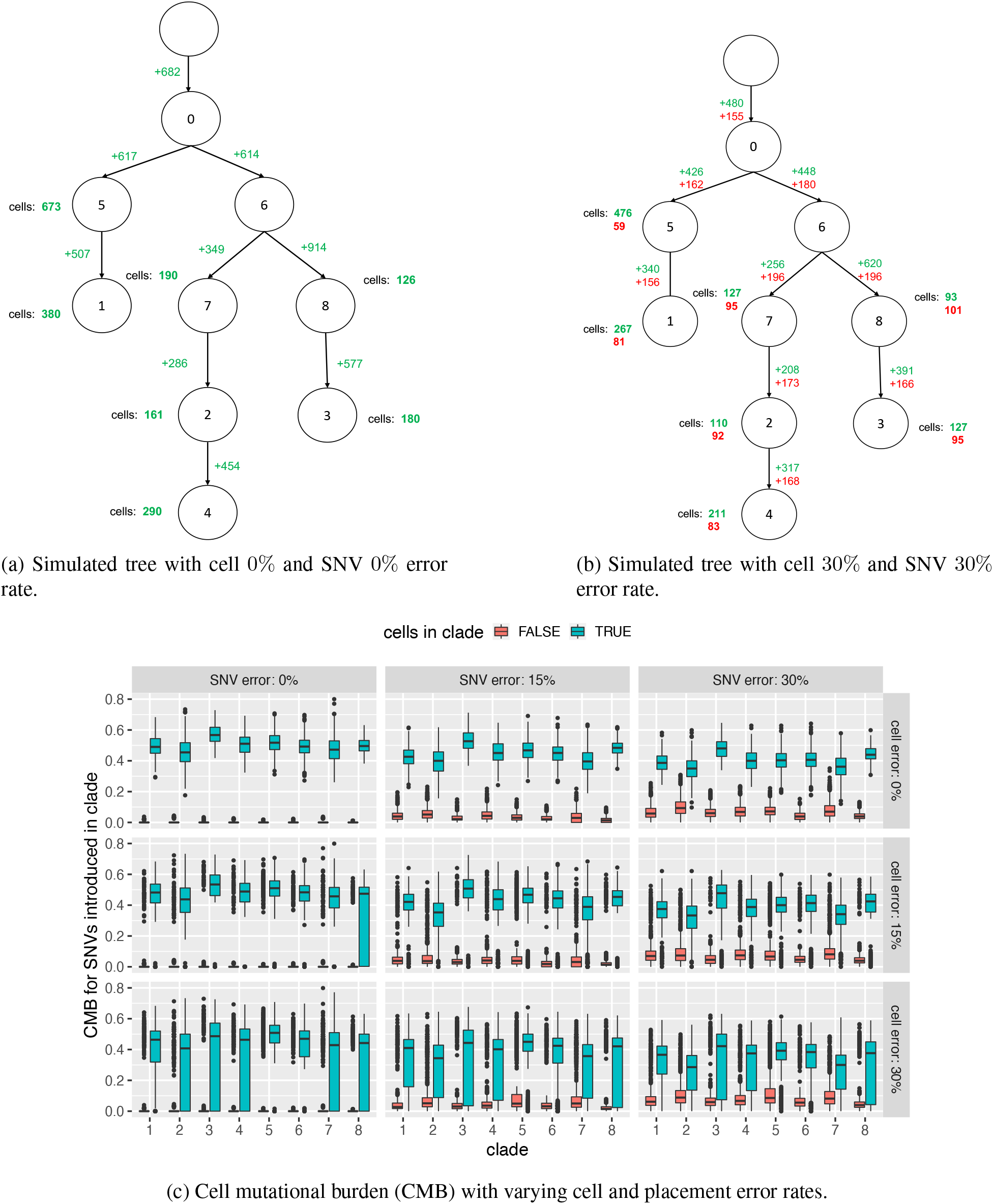
Analysis of cell mutational burden (CMB) on a simulated instance with varying error rates in {0%, 15%, 30%} for both cell and SNV placement. (a) The simulated ground truth tree with 0% cell placement error rate and 0% SNV placement error rate. (b) The simulated ground truth tree with 30% cell placement error rate and 30% SNV placement error rate. (c) Cell mutational burden (CMB) comparison between cells within (blue) and outside of (red) each clade in the inferred clonal tree at varying cell and SNV placement error rates.

#### B.2.4 Supplemental figures for high-grade serous ovarian cancer cells sequenced with DLP+

**Figure S14:**
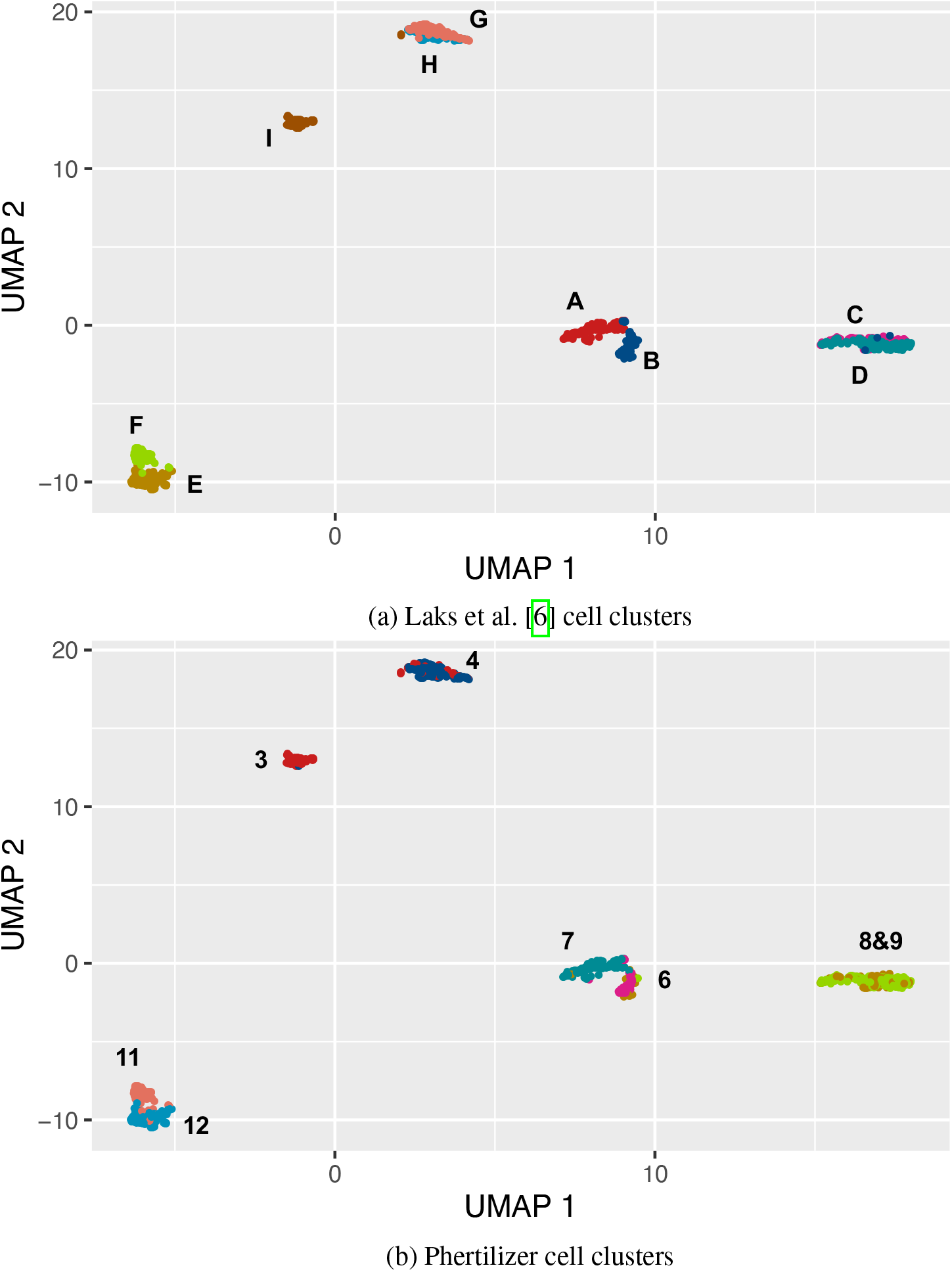
UMAP for the high-grade serous ovarian cancer patient depicted with inferred cell clusterings.

#### B.2.5 Processing the triple negative breast cancer tumors sequencing data from ACT

We merged reads from all cells into a single FASTQ file for each sample, aligned the merged reads to the Human reference genome hg19 (GRCh37) using bowtie2 (v.2.4.4), and sorted by samtools (v.1.15) forming pseudo-bulk samples. We then get a set of SNVs by running Mutect2 [38] in tumor-only mode on each pseudo-bulk sample, followed by FilterMutectCalls in GATK. We further select SNVs whose read depth is greater or equal to 11, VAF greater than 0.033, and have greater or equal to 4 variant reads in the pseudobulk sample. In order to match the bins in the original paper, we directly used the normalized read counts from ACT and find the low-dimensional embedding using UMAP [43].

#### B.2.6 Supplemental tables and figures for triple negative breast cancer tumors sequenced with ACT

- Table S1 provides a summary of the ACT data of all eight triple negative breast cancer tumors
- Fig. S15 shows a comparison of Phertilizer and Minussi et. al. [7] inferred cell clustering in the embedding space
- Fig. S16 shows the cell mutational burden (CMB) results for Baseline+SCITE inferred trees for breast tumors TN3 and TN5
- Fig. S17 shows Phertilizer supplemental results for breast tumor TN2
- Fig. S18 shows Phertilizer supplemental results for breast tumor TN4
- Fig. S19 shows Phertilizer supplemental results for breast tumor TN8

**Table S1:** Table depicting the tumor, number of cells and SNVs, the coverage, and the number of clones inferred by different methods for the eight analyzed breast cancer tumors.

| Tumor | Cells | SNVS | Coverage | Phertilizer clones | Minussi et al. super clones | Minussi et al. subclones | SBMclone clones |
| --- | --- | --- | --- | --- | --- | --- | --- |
| TN1 | 1100 | 13,934 | 0.031× | 6 | 4 | 17 | 1 |
| TN2 | 1024 | 8,474 | 0.039× | 6 | 4 | 15 | 1 |
| TN3 | 1101 | 13,064 | 0.020× | 1 | 4 | 9 | 1 |
| TN4 | 1307 | 14,333 | 0.022× | 5 | 4 | 22 | 1 |
| TN5 | 1238 | 10,271 | 0.020× | 1 | 4 | 7 | 1 |
| TN6 | 1205 | 11,827 | 0.017× | 1 | 4 | 15 | 1 |
| TN7 | 915 | 7,873 | 0.019× | 1 | 3 | 18 | 1 |
| TN8 | 1224 | 73,359 | 0.021× | 2 | 4 | 15 | 1 |

**Figure S15:**
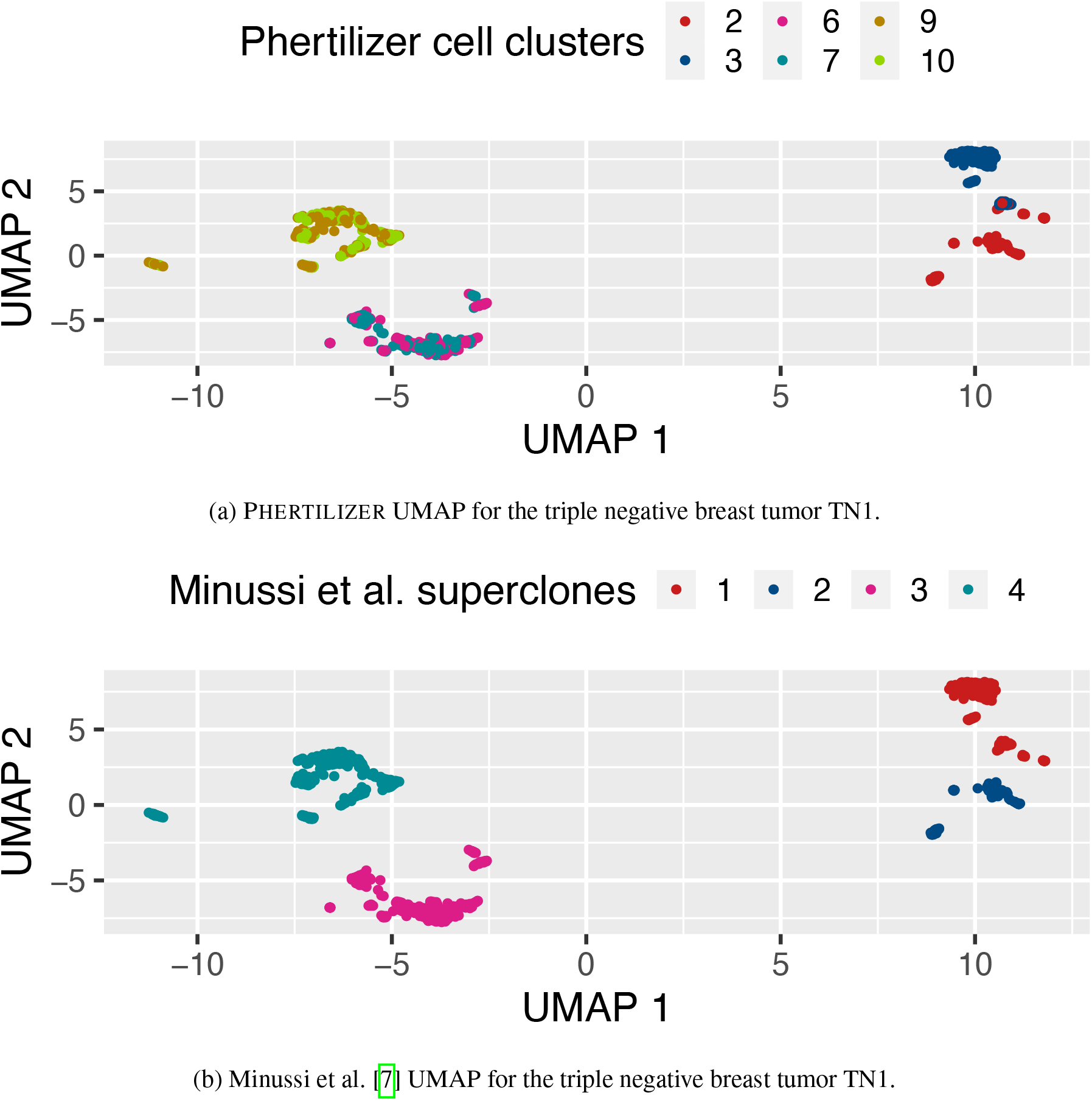
Cell clustering comparison for triple negative breast tumor TN1 in embedding space between Phertilizer and Minussi et al. [7].

**Figure S16:**
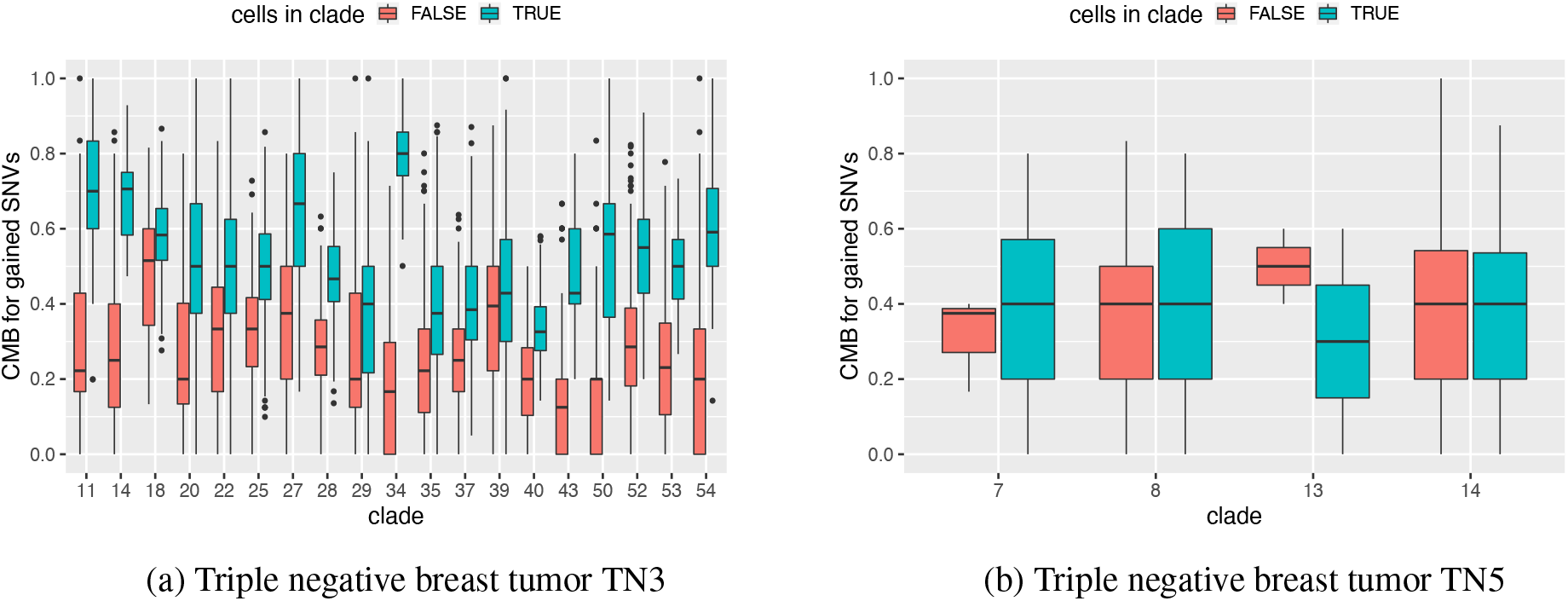
Cell mutational burden per clade in trees inferred by Baseline+SCITE for tumors TN3 and TN5.

**Figure S17:**
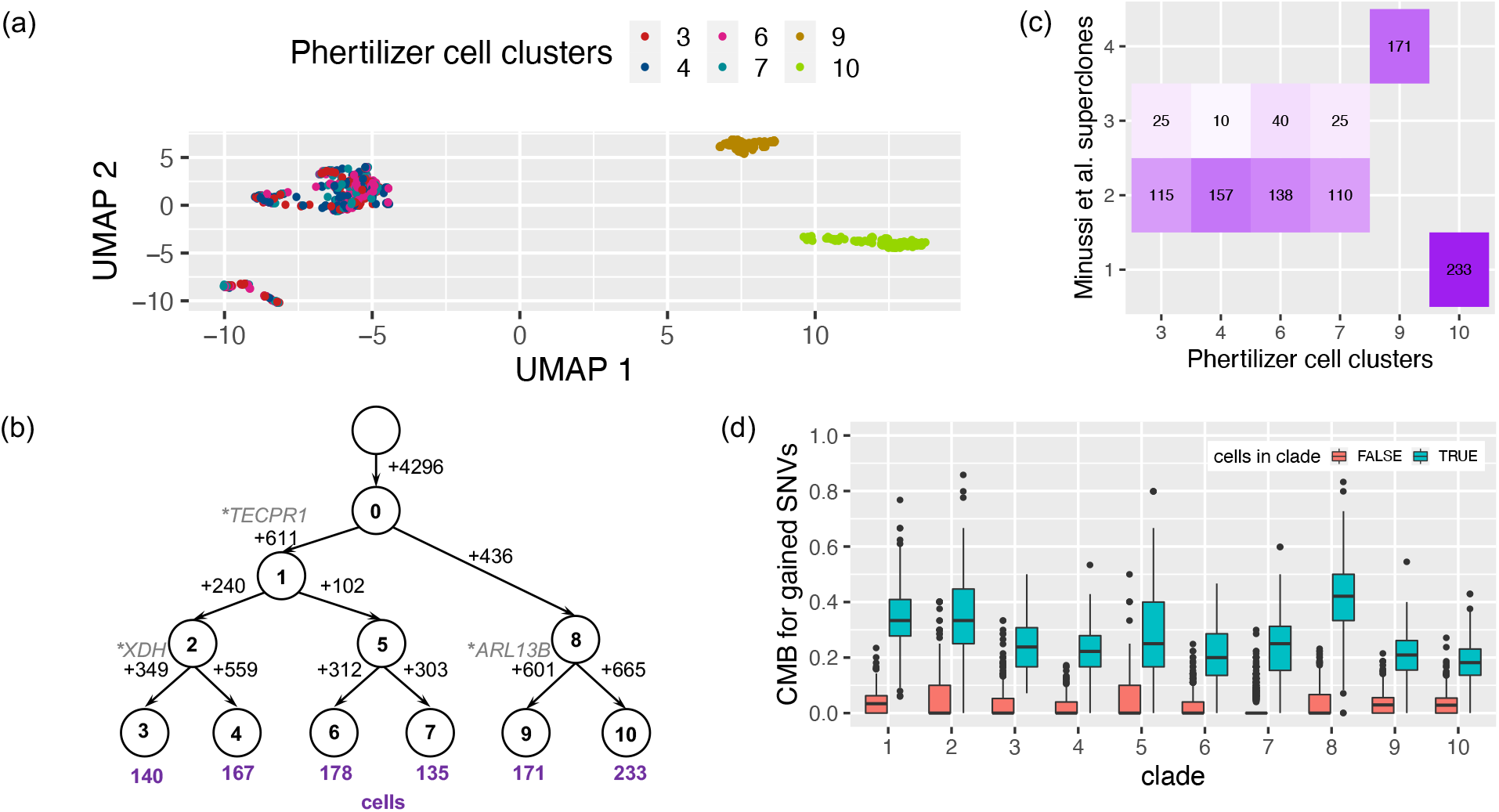
Phertilizer infers clonal tree for breast cancer tumor TN2. (a) UMAP for the triple negative breast tumor TN2. (b) The tree inferred by Phertilizer with numbers of SNVs labeled beside the edges, and numbers of cells labeled beneath the leaves. Cancer-related genes are labeled next to the SNVs (‘*’: stop-gain variant). (c) A mapping between Phertilizer’s cell clusters and the Minussi et al. [7] superclones. (d) The CMB distribution for the inferred clades for TN2.

**Figure S18:**
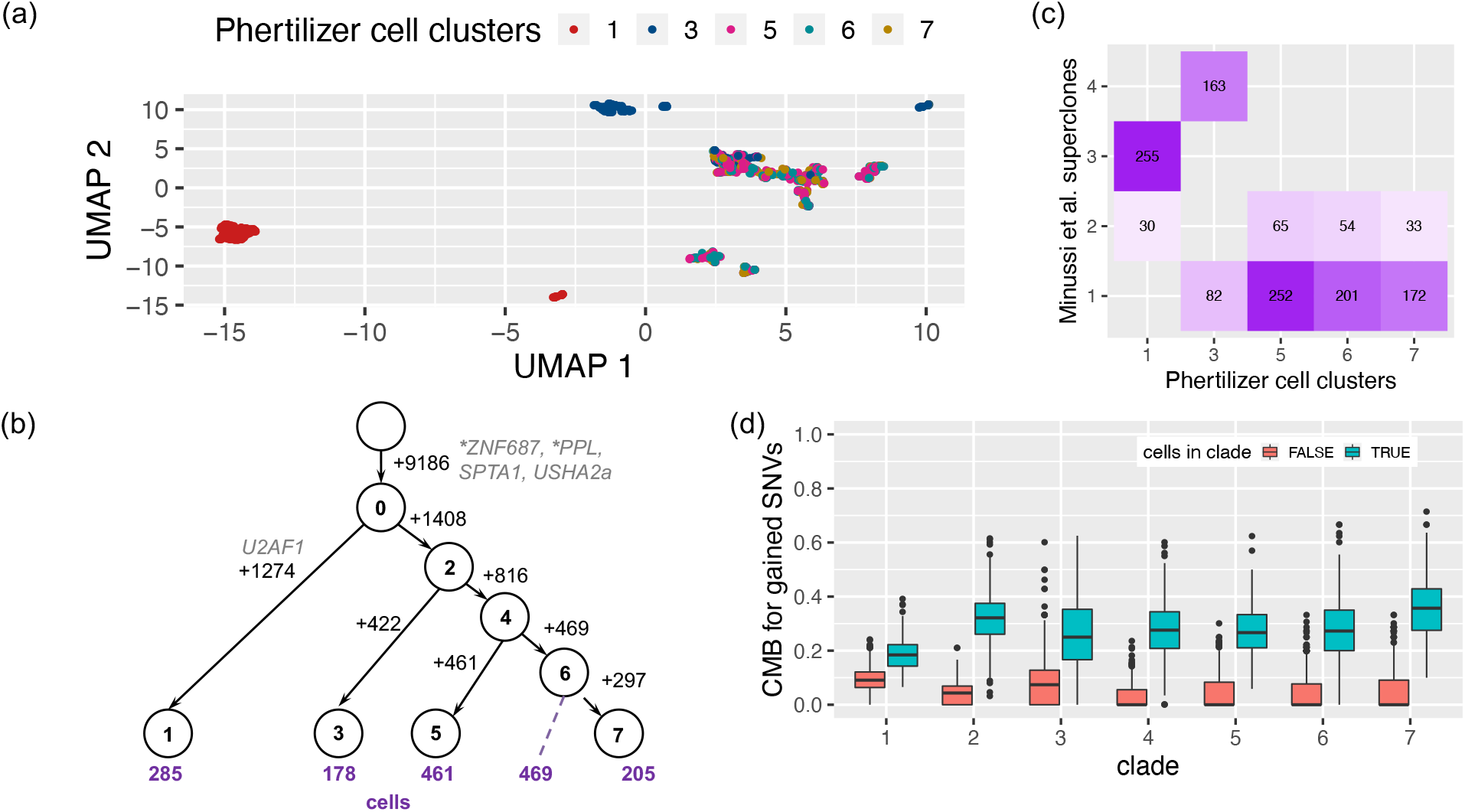
Phertilizer infers clonal tree for breast cancer tumor TN4. (a) UMAP for triple negative breast tumor TN4. (b) The tree inferred by Phertilizer with numbers of SNVs labeled beside the edges, and numbers of cells labeled beneath the leaves. Cancer-related genes are labeled next to the SNVs (‘*’: stop-gain variant). (c) A mapping between Phertilizer’s cell clusters and the Minussi et al. [7] superclones. (d) The CMB distribution for the inferred clades for TN4.

**Figure S19:**
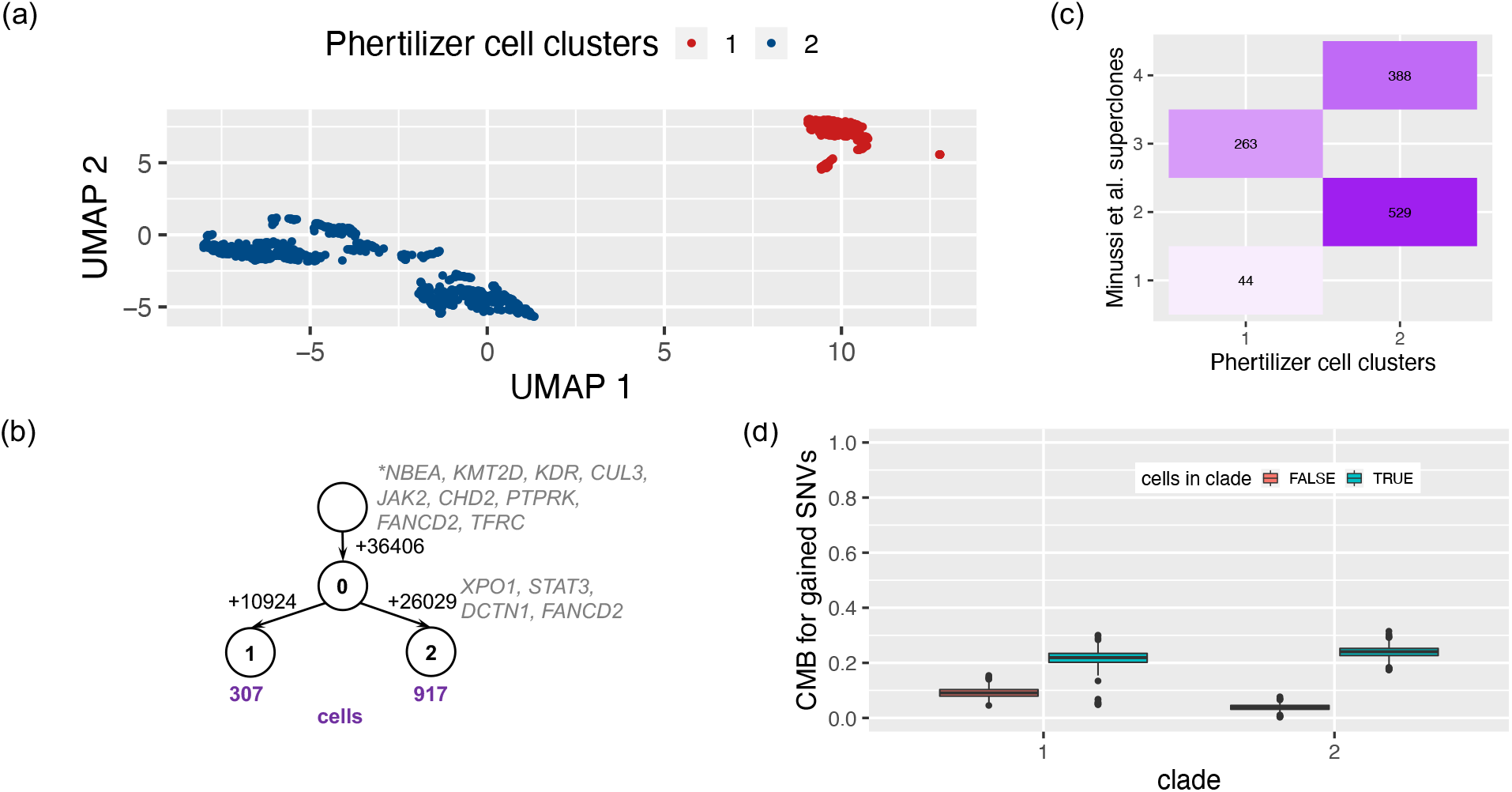
Phertilizer infers clonal tree for breast cancer tumor TN8. (a) UMAP for triple negative breast tumor TN8. (b) The tree inferred by Phertilizer with numbers of SNVs labeled beside the edges, and numbers of cells labeled beneath the leaves. Cancer-related genes are labeled next to the SNVs (‘*’: stop-gain variant). (c) A mapping between Phertilizer’s cell clusters and the Minussi et al. [7] superclones. (d) The CMB distribution for the inferred clades for TN8.

## References

[1] Nowell, P. C. The clonal evolution of tumor cell populations: Acquired genetic lability permits stepwise selection of variant sublines and underlies tumor progression. Science 194, 23–28 (1976).

[2] Morita, K. et al. Clonal evolution of acute myeloid leukemia revealed by high-throughput single-cell genomics. Nature Communications 11, 1–17 (2020).

[3] Baslan, T. et al. Novel insights into breast cancer copy number genetic heterogeneity revealed by single-cell genome sequencing. eLife 9, e51480 (2020).

[4] Kim, C. et al. Chemoresistance evolution in triple-negative breast cancer delineated by single-cell sequencing. Cell 173, 879–893 (2018).

[5] Miles, L. A. et al. Single-cell mutation analysis of clonal evolution in myeloid malignancies. Nature 587, 477–482 (2020).

[6] Laks, E. et al. Clonal decomposition and dna replication states defined by scaled single-cell genome sequencing. Cell 179, 1207–1221 (2019).

[7] Minussi, D. C. et al. Breast tumours maintain a reservoir of subclonal diversity during expansion. Nature 592, 302–308 (2021).

[8] Zahn, H. et al. Scalable whole-genome single-cell library preparation without preamplification. Nature Methods 14, 167–173 (2017).

[9] Pellegrino, M. et al. High-throughput single-cell DNA sequencing of acute myeloid leukemia tumors with droplet microfluidics. Genome Research 28, 1345–1352 (2018).

[10] Fu, X., Lei, H., Tao, Y. & Schwartz, R. Reconstructing tumor clonal lineage trees incorporating single-nucleotide variants, copy number alterations and structural variations. Bioinformatics 38, i125–i133 (2022).

[11] Kannan, J., Mathews, L., Wu, Z., Young, N. S. & Gao, S. CAISC: A software to integrate copy number variations and single nucleotide mutations for genetic heterogeneity profiling and subclone detection by single-cell rna sequencing. BMC bioinformatics 23, 1–17 (2022).

[12] Jahn, K., Kuipers, J. & Beerenwinkel, N. Tree inference for single-cell data. Genome Biology 17, 1–17 (2016).

[13] Malikic, S. et al. PhISCS: a combinatorial approach for subperfect tumor phylogeny reconstruction via integrative use of single-cell and bulk sequencing data. Genome Research 29, 1860–1877 (2019).

[14] El-Kebir, M. SPhyR: tumor phylogeny estimation from single-cell sequencing data under loss and error. Bioinformatics 34, i671–i679 (2018).

[15] Zafar, H., Navin, N., Chen, K. & Nakhleh, L. SiCloneFit: Bayesian inference of population structure, genotype, and phylogeny of tumor clones from single-cell genome sequencing data. Genome Research 29, 1847–1859 (2019).

[16] Roth, A. et al. Clonal genotype and population structure inference from single-cell tumor sequencing. Nature Methods 13, 573–576 (2016).

[17] Zaccaria, S. & Raphael, B. J. Characterizing allele-and haplotype-specific copy numbers in single cells with chisel. Nature Biotechnology 39, 207–214 (2021).

[18] Markowska, M. et al. CONET: Copy number event tree model of evolutionary tumor history for single-cell data. Genome Biology 23, 1–35 (2022).

[19] Liu, Y., Edrisi, M., Ogilvie, H. & Nakhleh, L. NestedBD: Bayesian inference of phylogenetic trees from single-cell DNA copy number profile data under a birth-death model. bioRxiv (2022).

[20] Wang, F. et al. MEDALT: single-cell copy number lineage tracing enabling gene discovery. Genome Biology 22, 1–22 (2021).

[21] Kaufmann, T. L. et al. MEDICC2: whole-genome doubling aware copy-number phylogenies for cancer evolution. Genome biology 23, 241 (2022).

[22] Kozlov, A., Alves, J. M., Stamatakis, A. & Posada, D. CellPhy: accurate and fast probabilistic inference of single-cell phylogenies from scDNA-seq data. Genome biology 23, 1–30 (2022).

[23] Kang, S. et al. SIEVE: joint inference of single-nucleotide variants and cell phylogeny from single-cell dna sequencing data. Genome Biology 23, 248 (2022).

[24] Chen, K., Moravec, J. C., Gavryushkin, A., Welch, D. & Drummond, A. J. Accounting for errors in data improves divergence time estimates in single-cell cancer evolution. Molecular biology and evolution 39, msac143 (2022).

[25] Milite, S., Bergamin, R., Patruno, L., Calonaci, N. & Caravagna, G. A Bayesian method to cluster single-cell RNA sequencing data using copy number alterations. Bioinformatics 38, 2512–2518 (2022).

[26] Zhou, Z., Xu, B., Minn, A. & Zhang, N. R. DENDRO: genetic heterogeneity profiling and subclone detection by single-cell RNA sequencing. Genome biology 21, 1–15 (2020).

[27] Satas, G., Zaccaria, S., Mon, G. & Raphael, B. J. SCARLET: single-cell tumor phylogeny inference with copy-number constrained mutation losses. Cell Systems 10, 323–332 (2020).

[28] Myers, M. A., Zaccaria, S. & Raphael, B. J. Identifying tumor clones in sparse single-cell mutation data. Bioinformatics 36, i186–i193 (2020).

[29] Rozhoňová, H. et al. SECEDO: SNV-based subclone detection using ultra-low coverage single-cell DNA sequencing. Bioinformatics 38, 4293–4300 (2022).

[30] Kimura, M. The number of heterozygous nucleotide sites maintained in a finite population due to steady flux of mutations. Genetics 61, 893 (1969).

[31] Davis, A., Gao, R. & Navin, N. Tumor evolution: Linear, branching, neutral or punctuated? Biochimica et Biophysica Acta (BBA)-Reviews on Cancer 1867, 151–161 (2017).

[32] Shi, J. & Malik, J. Normalized cuts and image segmentation. IEEE Transactions on pattern analysis and machine intelligence 22, 888–905 (2000).

[33] El-Kebir, M., Oesper, L., Acheson-Field, H. & Raphael, B. J. Reconstruction of clonal trees and tumor composition from multi-sample sequencing data. Bioinformatics 31, i62–i70 (2015).

[34] Dollo, L. Les lois de l’évolution. Bulletin de la Société belge de géologie, de paléontologie et d’hydrologie 7, 164–166 (1893).

[35] Cerami, E. et al. The cBio cancer genomics portal: an open platform for exploring multidimensional cancer genomics data. Cancer Discovery 2, 401–404 (2012).

[36] Gao, J. et al. Integrative analysis of complex cancer genomics and clinical profiles using the cBioPortal. Science Signaling 6, pl1–pl1 (2013).

[37] Sondka, Z. et al. The COSMIC Cancer Gene Census: describing genetic dysfunction across all human cancers. Nature Reviews Cancer 18, 696–705 (2018).

[38] Van der Auwera, G. A. & O’Connor, B. D. Genomics in the cloud: using Docker, GATK, and WDL in Terra (O’Reilly Media, 2020).

[39] Consortium, A. P. G. et al. AACR project GENIE: powering precision medicine through an international consortium. Cancer discovery 7, 818–831 (2017).

[40] Heravi-Moussavi, A. et al. Recurrent somatic DICER1 mutations in nonepithelial ovarian cancers. New England Journal of Medicine 366, 234–242 (2012).

[41] Sashittal, P., Zaccaria, S. & El-Kebir, M. Parsimonious clone tree integration in cancer. Algorithms for Molecular Biology 17, 1–14 (2022).

[42] El-Kebir, M., Morris, Q., Oesper, L. & Sahinalp, S. C. Emerging topics in cancer evolution. In PACIFIC SYMPOSIUM ON BIOCOMPUTING 2022, 397–401 (World Scientific, 2021).

[43] McInnes, L., Healy, J. & Melville, J. UMAP: Uniform manifold approximation and projection for dimension reduction. arXiv preprint arXiv:1802.03426 (2018).

[44] Weber, L. L., Sashittal, P. & El-Kebir, M. doubletD: detecting doublets in single-cell DNA sequencing data. Bioinformatics 37, i214–i221 (2021).

[45] De Bourcy, C. F. et al. A quantitative comparison of single-cell whole genome amplification methods. PloS one 9,e105585 (2014).

[46] Aggarwal, C. C., Hinneburg, A. & Keim, D. A. On the surprising behavior of distance metrics in high dimensional space. In International Conference on Database Theory, 420–434 (Springer, 2001).

[47] Kriegel, H.-P., Kröger, P. & Zimek, A. Clustering high-dimensional data: A survey on subspace clustering, pattern-based clustering, and correlation clustering. ACM Transactions on Knowledge Discovery from Data 3, 1–58 (2009).

[48] Pedregosa, F. et al. Scikit-learn: Machine learning in Python. Journal of Machine Learning Research 12, 2825–2830 (2011).

[49] Damle, A., Minden, V. & Ying, L. Simple, direct and efficient multi-way spectral clustering. Information and Inference: A Journal of the IMA 8, 181–203 (2019).

[50] Ankerst, M., Breunig, M. M., Kriegel, H.-P. & Sander, J. OPTICS: Ordering points to identify the clustering structure. ACM Sigmod record 28, 49–60 (1999).

[51] McLaren, W. et al. The Ensembl variant effect predictor. Genome biology 17, 1–14 (2016).

[52] Vaser, R., Adusumalli, S., Leng, S. N., Sikic, M. & Ng, P. C. Sift missense predictions for genomes. Nature protocols 11, 1–9 (2016).

[53] Adzhubei, I., Jordan, D. M. & Sunyaev, S. R. Predicting functional effect of human missense mutations using PolyPhen-2. Current protocols in human genetics 76, 7–20 (2013).

